# An Isogenic iPSC Model of LMNA p.R482W Recapitulates Adipocyte Dysfunction in Familial Partial Lipodystrophy Type 2

**DOI:** 10.64898/2026.10.29.760154

**Authors:** Aluet Borrego Alvarez, Will O. Wasielewski, McKinlee R.S. Gobble, Michael Grupe, Lina M. Ali, William Buchser, Lauren Boggs-Bailey, Dustin Baldridge, Jessica N. Maung, Akira Nishii, Ormond A. MacDougald, Elif A. Oral, Stephen I. Stone

## Abstract

Familial partial lipodystrophy type 2 (FPLD2) is a rare autosomal dominant disorder caused by pathogenic variants in *LMNA*, most commonly p.R482W, and is characterized by selective loss of subcutaneous adipose tissue, severe insulin resistance, and dyslipidemia. The cellular mechanisms underlying adipocyte dysfunction in FPLD2 remain incompletely understood, in part because existing *in vitro* models rely on patient-derived, non-isogenic cell lines or require non-physiologic transgene overexpression to achieve adipogenic differentiation. Here, we describe a CRISPR/Cas9-edited isogenic iPSC model of FPLD2 in which the heterozygous *LMNA* c.1444C>T (p.R482W) variant was introduced into a healthy donor-derived iPSC line. Using a transgene-free mesenchymal progenitor differentiation protocol, we generated *LMNA^p.R482W^* and isogenic control adipocytes and characterized their transcriptomic and functional phenotypes. Bulk RNA-sequencing demonstrated strong concordance in adipogenesis gene expression between genotypes during differentiation (Pearson ρ = 0.968), yet *LMNA^p.R482W^* adipocytes exhibited significant downregulation of key adipogenic, lipid-handling, and extracellular matrix genes compared to isogenic controls. Functionally, *LMNA^p.R482W^*adipocytes showed markedly impaired glucose uptake under both basal and insulin-stimulated conditions, consistent with severe insulin resistance. Seahorse extracellular flux analysis revealed a globally quiescent metabolic state with significantly reduced oxygen consumption rate (126.5 vs. 261.5 pmol/min, *p* < 0.0001) and extracellular acidification rate (27.6 vs. 46.7 mpH/min, *p* < 0.0001). To validate these findings in human tissue, gene set enrichment analysis results from the iPSC model were compared to single-nucleus RNA-sequencing data from subcutaneous white adipose tissue biopsies of individuals with FPLD2 carrying the *LMNA* p.R482W variant. Across 37 mutually significant pathways, 59.5% were concordant between the iPSC model and human adipose tissue, including the pathways most central to FPLD2 pathophysiology: adipogenesis, fatty acid metabolism, oxidative phosphorylation, and inflammatory response. These findings establish this isogenic iPSC system as a faithful cellular model of FPLD2 adipocyte biology and a platform for functional characterization of *LMNA* variants of uncertain significance.

**Graphical Abstract:** 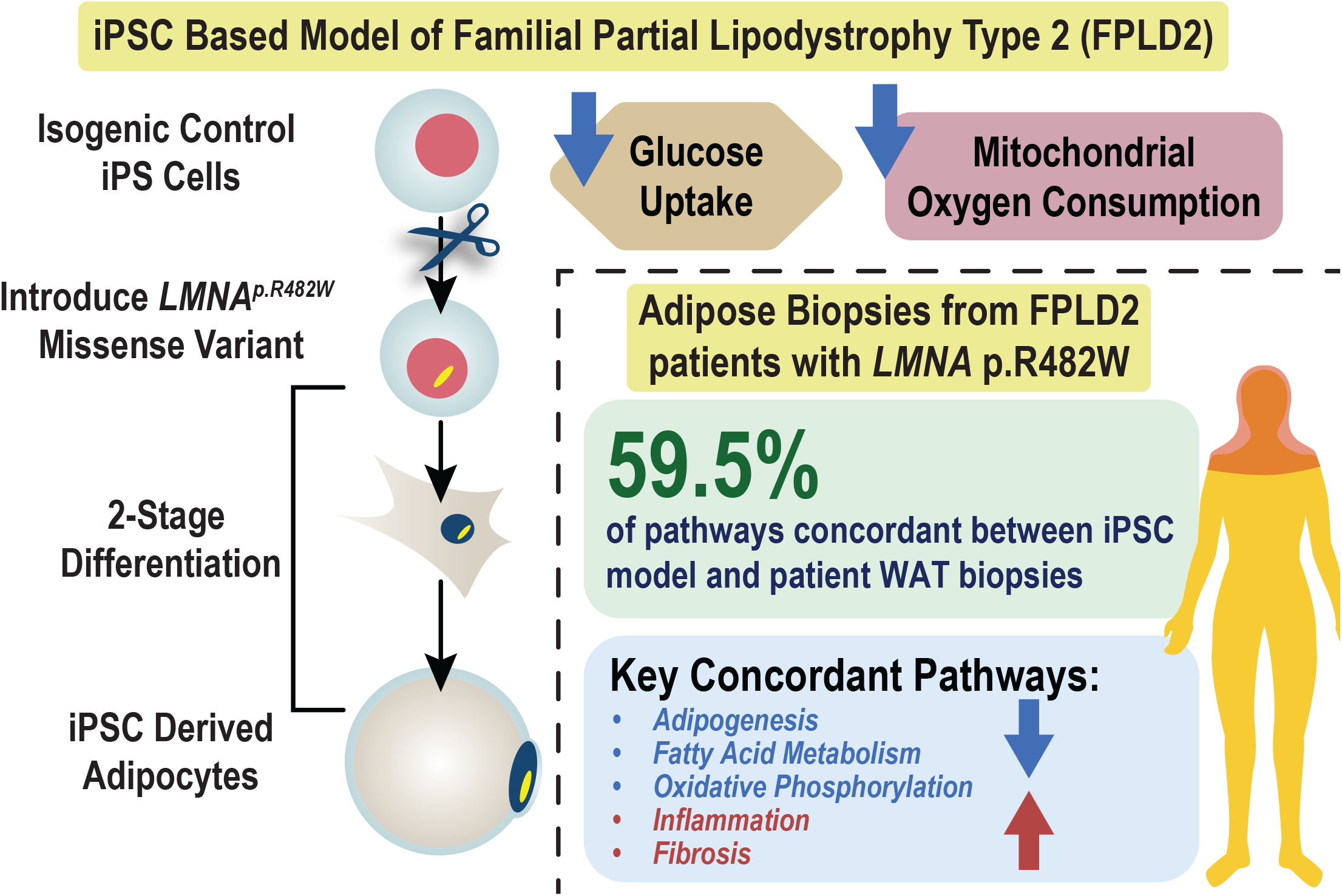

## Introduction

Adipose tissue is a central metabolic and endocrine organ, and alterations in its mass or function carry significant systemic consequences. Lipodystrophy syndromes are characterized by selective loss and redistribution of adipose tissue, resulting in metabolic complications that parallel those of severe obesity despite markedly reduced fat stores. Familial partial lipodystrophy type 2 (FPLD2, Dunnigan disease) is the most common monogenic lipodystrophy, caused by autosomal dominant missense variants in the *LMNA* gene and affecting approximately 1 in 20,000 individuals (1, 2). This loss and dysfunctional adipose tissue distribution drives severe metabolic complications including insulin resistance, diabetes mellitus, hypertriglyceridemia, and hepatic steatosis, all of which contribute substantially to morbidity (3). Available treatments address metabolic comorbidities rather than the primary adipocyte dysfunction, reflecting an incomplete understanding of the cellular mechanisms driving adipose loss in FPLD2.

*LMNA* encodes nuclear lamins A and C, intermediate filament proteins that form the inner nuclear lamina and regulate nuclear architecture, chromatin organization, and gene expression. The most common FPLD2-associated variant, R482W (c.1444C>T), disrupts a DNA-binding region in the globular tail of lamin A and is thought to cause transcriptional dysregulation of adipogenic programs, particularly those governed by PPARγ (4). Prior studies of FPLD2 adipose tissue identified suppression of lipid metabolic pathways and activation of inflammatory programs as key molecular features of the disease. Corsa and colleagues further demonstrated that lamin A/C is not required for adipogenesis per se, but that mature adipocytes lacking lamin A/C exhibit increased lipolytic sensitivity, pointing to a cell-autonomous defect in adipocyte maintenance rather than differentiation (5). Maung and colleagues recently provided the most comprehensive human tissue characterization of FPLD2 to date, profiling subcutaneous white adipose tissue biopsies from individuals with pathogenic *LMNA* variants enrolled in the Longitudinal Evaluation of Adiposity Distribution and Adipocyte Biology in Children with Lipodystrophy (LEAD-ABC) study, and integrating these findings with an adipocyte-specific *Lmna*-knockout mouse model (6). That work established conserved suppression of lipid metabolism and fatty acid oxidation alongside inflammatory pathway activation as hallmark transcriptomic signatures of FPLD2 and identified adipocyte degeneration as a primary mechanism of fat loss. However, the intrinsic molecular mechanisms driving these transcriptomic signatures in adipocytes carrying the R482W variant specifically have not been directly interrogated.

Human cellular models are essential tools for mechanistic investigation of adipocyte-intrinsic disease. Patient-derived fibroblasts and preadipocyte cultures preserve the human genetic context but are limited in scalability, differentiation potential, and ability to model adipogenic commitment. Induced pluripotent stem cell (iPSC) technology overcomes many of these limitations by providing a renewable, genetically defined source of human cells that can be differentiated into mature adipocytes (7, 8). Friesen and Cowan provided an early proof of concept by generating iPSC-derived adipocytes from two patients with FPLD2 and demonstrating reduced adipogenic marker expression and impaired insulin signaling (9). However, that study compared non-isogenic patient-derived lines to unrelated healthy controls, making it difficult to separate variant-specific effects from inter-individual genetic background variation. Furthermore, lentiviral *PPARG2* overexpression was required to drive adipogenic commitment, which may obscure the endogenous transcriptional response to the *LMNA* variant. No prior iPSC model of FPLD2 has been validated against patient-derived tissue data, leaving the translational fidelity of these *in vitro* systems uncertain. Beyond mechanistic research, there is also an unmet clinical need for a platform to functionally characterize *LMNA* variants of uncertain significance in patients with atypical presentations of partial lipodystrophy, a growing and underdiagnosed population for whom genetic testing often yields ambiguous results (2).

To address these gaps, we generated an isogenic iPSC model of FPLD2 by introducing the pathogenic *LMNA* c.1444C>T (p.R482W) variant into a single isogenic control iPSC line using CRISPR/Cas9 genome editing, producing two independent heterozygous clones for comparison against the unmodified parental line. Cells were differentiated to white adipocytes through a commercially defined, transgene-free mesenchymal progenitor protocol and subjected to bulk RNA-sequencing, 2-deoxyglucose uptake assays, and mitochondrial stress testing. To establish translational validity, transcriptomic results were cross-validated against single nuclei RNA sequencing (snRNA-seq) data from the R482W-specific subset of the LEAD-ABC cohort reported by Maung and colleagues, enabling direct comparison between the *in vitro* model and human adipose tissue from individuals carrying the identical variant (6). We demonstrate that *LMNA^p.R482W^* adipocytes faithfully recapitulate the transcriptomic, metabolic, and inflammatory signatures of human FPLD2 adipose tissue, resolve discrepancies between prior *in vitro* findings, and establish a validated human cellular platform for mechanistic investigation, therapeutic screening, and the functional evaluation of *LMNA* variants of uncertain significance.

### iPSC line and CRISPR/Cas9 genome editing

AN1.1 induced pluripotent stem cells (iPSCs) were previously reprogrammed from renal epithelial cells isolated from a healthy adult woman (10). The pathogenic lamin A/C variant (*LMNA*; NM_170707.4:c.1444C>T; p.(R482W)) was introduced into this isogenic control iPSC line using CRISPR/Cas9 genome editing. The following guide RNAs (gRNAs) were used:

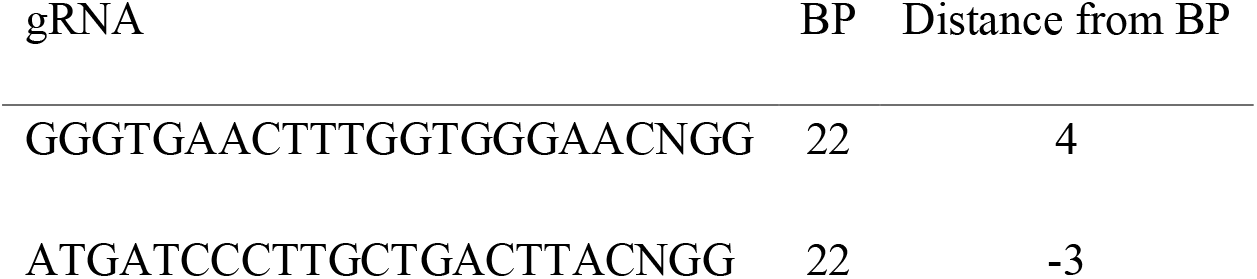

Two clones were successfully edited to express the heterozygous *LMNA* c.1444C>T variant and were labeled Het 2A4 and Het 4A5. Next-generation sequencing was used to confirm the intended edit without evidence of off-target insertions or deletions. Quality control included karyotyping and short tandem repeat (STR) profiling (11), both of which returned normal results. Proper iPSC morphology was confirmed by light microscopy prior to downstream differentiation.

### iPSC culture and mesenchymal progenitor cell differentiation

iPSCs were maintained in mTeSR™ Plus medium (StemCell Technologies, #100-0276). To initiate mesenchymal differentiation, isogenic control and *LMNA^p.R482W^* iPSCs were seeded and cultured using the STEMdiff™ Mesenchymal Progenitor Kit (StemCell Technologies, #05240). Cells were considered ready for induction when most colonies demonstrated dense centers and compact morphology. After 2 days of complete media changes in mTeSR™ Plus media, the media was aspirated and replaced with STEMdiff™-ACF Mesenchymal Induction Medium (StemCell Technologies, #05241) to induce early mesenchymal progenitor cells. After 1-3 additional full media changes, cells were transitioned to MesenCult™-ACF Plus Medium (StemCell Technologies, #05446) for two passages. Mesenchymal progenitor cell (MPC) identity was confirmed by the characteristic elongated, spindle-shaped morphology assessed by light microscopy.

Mesenchymal progenitor cells were expanded for four passages prior to the initiation of adipogenic differentiation. To induce adipogenesis, medium was switched to MesenCult™ Adipogenic Differentiation Kit (Human; StemCell Technologies, #05412). Medium was replaced every 2-3 days over a 21-day differentiation period. Successful adipogenic differentiation was confirmed by accumulation of cytoplasmic lipid droplets that coalesced into a multilocular pattern resembling mature adipocytes.

### Nile Red staining

Lipid droplet formation was confirmed by Nile Red staining. Media containing Nile Red (1:1000; Sigma-Aldrich; #298395) and Hoechst 33342 (1:3000; Sigma-Aldrich; #TA9H97BAECD2) was added slowly to each well. After 20 minutes, the stain was removed by gently aspirating the media and washed once with 100 µL PBS. 100 µL fresh media was added back to the wells prior to imaging. Cells were imaged on the Molecular Devices InCell6500 microscope at 20X magnification with 1x1 Binning, and Laser Autofocus at 6% power.

### Bulk RNA-sequencing

RNA was extracted from MPCs and adipocytes cultured in individual wells of 6-well plates. Briefly, cells were washed with PBS, 1 mL of TRIzol (Invitrogen) was added, cells were scraped and transferred to Eppendorf tubes. 200 µL of chloroform was added, followed by centrifugation and retention of the aqueous layer. RNA was precipitated with isopropanol and centrifugation, followed by washing with 75% ethanol. Column-based RNA cleanup was performed using the Monarch Spin RNA Cleanup Kit (NEB). Samples were submitted to Novogene (Sacramento, CA) for mRNA library preparation with polyA enrichment and NovaSeq paired end 150 bp sequencing, with a target of 6 Gb of raw data per sample. Raw reads were aligned to the human reference genome (GRCh38) using HISAT2 (v2.0.5), and gene-level expression was quantified as fragments per kilobase per million mapped reads (FPKM) using featureCounts (v1.5.0-p3). Differential expression analysis was performed using DESeq2 (v1.20.0), with statistical significance defined as a false discovery rate (FDR) of 0.05 or less and an absolute log_2_ fold change of 1 or greater. Genes were ranked by Wald test statistic and used as input for gene set enrichment analysis (GSEA) using the fgsea R package (v.1.34.2) (12), with the MSigDB Hallmark gene set collection (12, 13) in R (4.5.1). Pearson correlation analysis was used to assess concordance in gene-level expression changes between isogenic control and *LMNA^p.R482W^*cell lines across differentiation, as well as concordance within the adipogenesis gene set specifically.

### 2-Deoxyglucose uptake assay

Insulin sensitivity was assessed in day-21 adipocytes using a fluorometric 2-deoxyglucose (2-DG) uptake assay (Promega; Glucose Uptake-Glo™ Assay; #J1342). 10,000 cells/well were plated in a 96-well tissue culture plate (BRANDplates^®^, #781970) for 2 days to reach 90% confluency. Prior to beginning the 2-DG assay, the cells were serum starved for 16 hours. Cell Viability Assay (CellTiter-Glo^®^ Assay; # G9241) was performed in conjunction with the 2-DG assay to confirm equal confluency. Subsequently, the cells were incubated (37 ℃; 5% CO2) for one hour with recombinant human insulin (Humulin R; 0002-8215-01) at concentrations of 0, 10, 100, and 1000 nM. Following insulin treatment, cells were incubated with 2-DG (1 mM) for 10 minutes at room temperature. Relative light units (RLU) were measured using a TECAN Infinite^®^ M Plex multimode microplate reader. Statistical comparisons between isogenic control and *LMNA^p.R482W^* cells were performed at each insulin concentration using a two-tailed unpaired *t*-test.

### Seahorse XFe96 assay

Cellular respiration was assessed using the Seahorse XFe96 Extracellular Flux Analyzer (Agilent Technologies). Day-21 adipocytes were seeded in Seahorse XFe96 cell culture microplates at a density of 10,000 cells per well 5 days prior to the assay. Cells were equilibrated in Seahorse XF base medium (Agilent Technologies; #103335-100) supplemented with 10 mM glucose, 1 mM pyruvate, 2 mM glutamine at 37 °C in a non-CO_2_ incubator for one hour prior to measurement. The Mito Stress Test (Agilent Technologies; #103015-100) was performed using sequential injections of oligomycin (1 µM), carbonyl cyanide 4-(trifluoromethoxy) phenylhydrazone (FCCP; 0.5 µM), and a combination of rotenone and antimycin A (0.5 µM). Oxygen consumption rate (OCR) and extracellular acidification rate (ECAR) were measured at baseline and following each injection. Values were normalized to cell number by delivering Hoechst 33422 (1:3000; Sigma-Aldrich; #TA9H97BAECD2) together with Rotenone/Antimycin via the third port to live cells. Images were taken at the end of the Seahorse Assay using a BioTek Cytation 5 Cell Imaging Multimode Reader. Energy phenotype mapping was performed by plotting ECAR against OCR under basal and maximally stimulated conditions for each cell line. Statistical comparisons of OCR and ECAR were made using a two-tailed unpaired *t*-test.

Single-nucleus RNA-sequencing (snRNA-seq) data from subcutaneous white adipose tissue (WAT) biopsies were obtained from a subset of participants enrolled in the Longitudinal Evaluation of Adiposity Distribution and Adipocyte Biology in Children with Lipodystrophy (LEAD-ABC) study. Participant recruitment, informed consent, and biopsy collection were conducted under protocols approved by the relevant institutional review board, as described in Maung et *al.* (6). For the present analysis, biopsies from 3 individuals carrying the *LMNA* p.R482W variant and 5 unaffected control individuals were included. Biopsies were collected from two anatomical sites: the upper neck (a region of lipohypertrophy in FPLD2) and the abdomen (a region of lipoatrophy), to enable comparison across metabolically distinct adipose depots.

Probes were hybridized, and samples were pooled and processed per manufacturer instructions for the 10x Genomics Chromium Fixed RNA Profiling Reagent Kits for Multiplexed Samples (#1000568). Library quality was assessed using the LabChip GXII HT (PerkinElmer) and quantified by Qubit (Thermo Fisher Scientific). Pooled libraries underwent paired-end sequencing on an Illumina NovaSeq XPlus. BCL Convert Software (Illumina) generated demultiplexed FASTQ files, and the Cell Ranger Pipeline (10x Genomics) was used for alignment and count matrix generation. Seurat v5.4.0 was used for filtering, normalization, dimensional reduction, clustering, gene expression visualization, and differential expression analysis. Cells were excluded if they had <1,000 unique molecular identifiers (UMIs), <500 detected genes, >5% mitochondrial transcripts, or log_10_ genes per UMI < 0.80; genes were excluded if detected in <10 cells. SCTransform was used for normalization and variance stabilization, treating mitochondrial mapping percentage as a covariate. To integrate and correct for batch effects, Harmony was applied to the SCTransformed data via RunHarmony. Resulting embeddings were used for downstream dimensional reduction and clustering (6).

Seven major cell populations were identified: adipocytes, adipose stem and progenitor cells (ASPCs), macrophages, endothelial cells, T cells, pericytes, and lymphatic endothelial cells (LECs). Cell type identity was assigned based on expression of canonical marker genes (*ADIPOQ*, *PDGFRA*, *MAFB*, *JAM2*, *IL7R*, *STEAP4*, *PROX1*) as previously described (14). Cell populations were visualized using Uniform Manifold Approximation and Projection (UMAP). GSEA was performed on the adipocyte cluster comparing *LMNA^p.R482W^* individuals to controls using the MSigDB Hallmark gene set collection (13).

To assess the degree to which the *in vitro* iPSC model recapitulates transcriptomic signatures observed in human FPLD2 adipose tissue, normalized enrichment scores (NES) from the iPSC bulk RNA-seq GSEA were compared against NES from the snRNA-seq GSEA of adipocyte clusters from upper neck and abdominal biopsies. Only gene sets achieving statistical significance (FDR < 0.05) in both datasets being compared were included in each pairwise analysis. Pathways were classified as concordant (NES in the same direction in both datasets), partially concordant (significant in one dataset only), or discordant (NES in opposite directions). Pearson and Spearman correlation coefficients were calculated for each pairwise comparison. For the three-way comparison (iPSC, upper neck, and abdomen), concordance was determined by requiring agreement in direction across all three datasets simultaneously.

### Statistical analysis

Statistical analyses were performed using GraphPad Prism (version 11) and R (version 4.5.1). For 2-DG uptake and Seahorse assays, differences between isogenic control and *LMNA^p.R482W^* cells were evaluated using a two-tailed unpaired *t*-test. For bulk RNA-seq differential expression, statistical significance was defined as an adjusted p-value (FDR) of 0.05 or less and an absolute log_2_ fold change of 1 or greater, after correction for multiple testing using the Benjamini-Hochberg method. For GSEA, statistical significance was defined as FDR < 0.25 as recommended by the GSEA software documentation (13). Pearson and Spearman correlation coefficients were used to assess concordance between transcriptomic datasets as described above. For all comparisons, a *p* value less than 0.05 was considered statistically significant unless otherwise noted. Because the differences between the two edited iPSC clones (Het 2A4 and Het 4A5) were negligible across functional assays, results from both lines were combined in the main figures; individual line data are presented separately in the supplemental figures.

iPSCs were derived from a female donor, which is especially relevant for modeling FPLD2, as the disease shows greater phenotypic penetrance in females. This sex bias is thought to reflect hormonal influences on adipose tissue biology (15). Findings based on *in vitro* work are nonetheless expected to have relevance to both sexes. For biopsy-derived data, both sexes were represented; however, sex was not included as a covariate due to the extreme rarity of this condition and the resulting small sample size (*n* = 8).

### Data availability

Raw and processed bulk RNA-sequencing data generated in this study will be deposited in the NCBI Gene Expression Omnibus (GEO) upon publication in a peer reviewed journal. The single-nucleus RNA-sequencing data from the LEAD-ABC cohort used for cross-platform validation are available through the repository associated with Maung *et al*.(6), under accession number GSE310542. Code used for differential expression analysis, GSEA, and concordance analysis will also be made available in a GitHub repository upon publication. Additional data supporting the findings of this study are available from the corresponding author upon request.

### Genome editing to generate heterozygous *LMNA^p.R482W^* lines

A isogenic control induced pluripotent stem cell (iPSC) line was previously reprogrammed from the renal epithelial cells of a healthy adult woman. To develop an iPSC line modeling FPLD2, the *LMNA* (c.1444C>T; p.R482W) variant was introduced into AN1.1 cells using CRISPR/Cas9 genome editing (**Fig. 1A**). Two heterozygous lines (Het 2A4 and Het 4A5) with the c.1444C>T variant were generated. The individual results for functional testing of each of these lines are plotted separately in the supplemental data figures (**Fig. S1**). However, the differences between the lines were mostly negligible, therefore results of both lines are combined in the figures (denoted as *LMNA^p.R482W^*). After genome editing, iPSCs underwent quality control steps including karyotyping, short-tandem repeat profiling (11), and next-generation sequencing.

**Figure 1.**
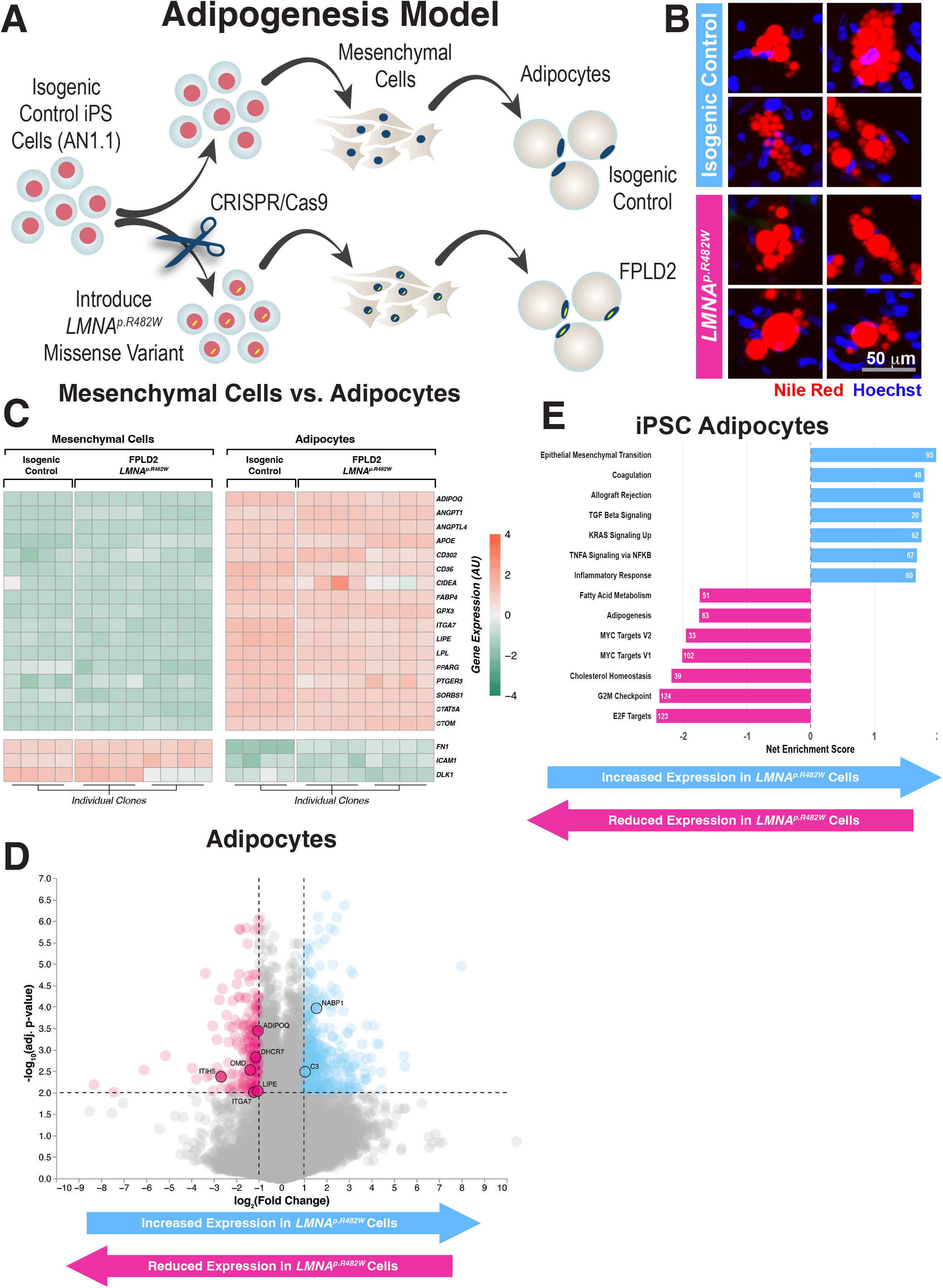
Generation and transcriptomic characterization of isogenic *LMNA^p.R482W^* iPSC-derived adipocytes. **(A)** Schematic of the iPSC adipogenesis model. The isogenic control AN1.1 iPSC line was edited by CRISPR/Cas9 to introduce the heterozygous *LMNA* c.1444C>T (p.R482W) variant. Both isogenic control and *LMNA^p.R482W^* iPSCs were differentiated through a mesenchymal progenitor cell (MPC) intermediate to mature adipocytes using a transgene-free protocol. **(B)** Representative fluorescence microscopy images of isogenic control (top) and *LMNA^p.R482W^*(bottom) iPSC-derived adipocytes at day 21 of differentiation. Cells were stained with Nile Red (lipid droplets, red) and Hoechst (nuclei, blue). Scale bar, (50) µm. **(C)** Heatmap of differential gene expression in each cell line before and after adipogenic differentiation (adipocytes vs. MPCs). At the top are the MSigDB Hallmark Adipogenesis gene set, followed by three mesenchymal cell markers. **(D)** Volcano plot of differential gene expression between isogenic control and *LMNA^p.R482W^* adipocytes. Pink points indicate genes with reduced expression in *LMNA^p.R482W^* adipocytes; blue points indicate genes with increased expression in *LMNA^p.R482W^* adipocytes. Labeled genes are members of the MSigDB Hallmark Adipogenesis gene set that achieved statistical significance. Dashed lines indicate log_2_ fold change thresholds of ±1 and adjusted p-value threshold of 0.05. **(E)** Gene set enrichment analysis (GSEA) bar chart for iPSC-derived adipocytes comparing *LMNA^p.R482W^* to isogenic control. Blue bars (positive net enrichment score, NES) indicate pathways enriched in *LMNA^p.R482W^* adipocytes; pink bars (negative NES) indicate pathways with reduced expression in *LMNA^p.R482W^* adipocytes. Numbers indicate leading edge gene counts.

### *LMNA^p.R482W^* iPSCs differentiate to mesenchymal progenitor cells with normal morphology

Isogenic control and *LMNA^p.R482W^* cells were differentiated to mesenchymal progenitor cells (MPCs) according to manufacturer instructions (**Fig. 1A**). MPCs were generated after 2 passages in MesenCult™-ACF Plus Medium. On light microscopy, the MPCs exhibited an elongated spindle-shaped morphology confirming MPC state.

### *LMNA^p.R482W^* mesenchymal progenitors form lipid-laden adipocytes

MPCs were expanded for 4 additional passages prior to starting adipogenic differentiation. To induce adipogenesis, the media was changed to the MesenCult™ Adipogenic Differentiation Kit (Human). Media was replaced every 2-3 days. Over 21 days of adipogenic differentiation, the MPCs accumulated cytoplasmic lipid droplets. Towards the end of the 21-day period, lipid droplets coalesced and the cells resembled mature adipocytes with multilocular lipid droplets (**Fig. 1A**). Mature-like adipocytes were confirmed via staining with Nile Red (**Fig. 1B**).

### *LMNA^p.R482W^* adipocytes exhibit decreased adipogenic gene expression

To interrogate the differentiation of mesenchymal cells to adipocytes, isogenic control and *LMNA^p.R482W^* cells underwent bulk RNA-sequencing before and after adipogenic differentiation. When exploring differential expression between the mesenchymal and adipocyte lines, we analyzed a subset of key marker genes involved in adipogenesis (13). As expected, several adipogenesis markers were upregulated in isogenic control adipocytes compared to their mesenchymal precursors *FABP4*, *GPX3*, *ITGA7*, *LIPE*, *LPL*, *PPARG*, *PTGER3*, *SORBS1*, *STAT5A*, and *STOM* were significantly increased in the adipocytes post-differentiation (**Fig. S2**, **Table S1**).

To determine whether isogenic control and *LMNA^p.R482W^* cells mounted a similar transcriptional response to adipogenic differentiation, we compared genome-wide fold-change profiles (MPC vs. adipocyte) between genotypes. The two profiles were strongly correlated (Pearson ρ = 0.700), indicating broadly parallel differentiation-associated gene expression changes. Correlation was even stronger when restricted to genes in the Hallmark Adipogenesis gene set (Pearson ρ = 0.968) *LMNA^p.R482W^* adipocytes similarly upregulated many Hallmark Adipogenesis genes during differentiation, with a pattern largely overlapping that of isogenic control cells (**Fig. S2**). However, two genes that were significantly upregulated in isogenic control adipocytes, *CIDEA* and *LPL*, did not reach significance in *LMNA^p.R482W^* cells, while *FZD4* was uniquely upregulated in *LMNA^p.R482W^* cells (**Table S2**).

To study effects of the pathogenic *LMNA^p.R482W^* variant, we compared bulk RNA-sequencing data results within the mesenchymal cells across genotypes (**Fig. S3A**). In the mesenchymal cells, only the expression of *ITIH5* was significantly decreased in *LMNA^p.R482W^* cells compared to isogenic control (*LMNA^p.R482W^* 1026.16 FPKM; isogenic control 3887.27 FPKM; *p* = 2.91 × 10^−5^) (**Fig. S3B**). In contrast, the adipocytes demonstrated several differentially expressed adipogenesis genes (**Fig. 1D**). Of the Hallmark Adipogenesis genes, *ADIPOQ*, *ITIH5*, *DHCR7*, *LIPE*, *ITGA7*, and *OMD* were significantly decreased in *LMNA^p.R482W^* cells, whereas *NABP1* and *C3* were significantly decreased in the isogenic control cells (**Fig. S3C** and **Table S3**).

Gene set enrichment analysis (GSEA) was utilized to identify the top enriched and depleted pathways in the iPSC-derived adipocytes (comparing *LMNA^p.R482W^* to isogenic control). Epithelial mesenchymal transition, coagulation, allograft rejection, TGF-beta signaling, KRAS signaling up, TNFA signaling via NFκB, and inflammatory response were enriched in the isogenic control cells. Conversely, the *LMNA^p.R482W^* cells demonstrated depletion of E2F targets, G2M checkpoint, cholesterol homeostasis, Myc targets V1, Myc targets V2, adipogenesis, and fatty acid metabolism (**Fig. 1E**).

### *LMNA^p.R482W^* adipocytes demonstrate severe insulin resistance under basal and stimulated conditions

To examine insulin sensitivity, isogenic control and *LMNA^p.R482W^* adipocytes were treated with increasing concentrations of recombinant human insulin, and glucose uptake was assessed by measuring 2-deoxyglucose (2-DG) uptake. Under basal conditions (0 nM insulin), *LMNA^p.R482W^* cells demonstrated significantly lower 2-DG uptake compared to isogenic control (WT 1.91 ± 0.48 × 10^6^ RLU vs *LMNA^p.R482W^* 0.87 ± 0.09 × 10^6^ RLU; *p* = 0.0001). Insulin-stimulated 2-DG uptake remained significantly attenuated in *LMNA^p.R482W^* cells at all tested concentrations, including 10 nM (WT 2.31 ± 0.98 × 10^6^ vs mutant 1.13 ± 0.24 × 10^6^ RLU; *p* = 0.007), 100 nM (WT 2.59 ± 1.15 × 10^6^ vs mutant 1.26 ± 0.24 × 10^6^ RLU; *p* = 0.009), and 1000 nM insulin (WT 4.18 ± 1.29 × 10^6^ vs mutant 1.78 ± 0.47 × 10^6^ RLU; *p* = 0.0006) (**Fig. 2A** and **Fig. S1A**). No significant difference in cell viability was detected between isogenic control and *LMNA^p.R482W^* wells prior to the assay, confirming that differences in 2-DG uptake reflect insulin resistance rather than differences in cell number/confluency. Overall, *LMNA^p.R482W^*adipocytes exhibited reduced basal glucose uptake and blunted insulin responsiveness.

**Figure 2.**
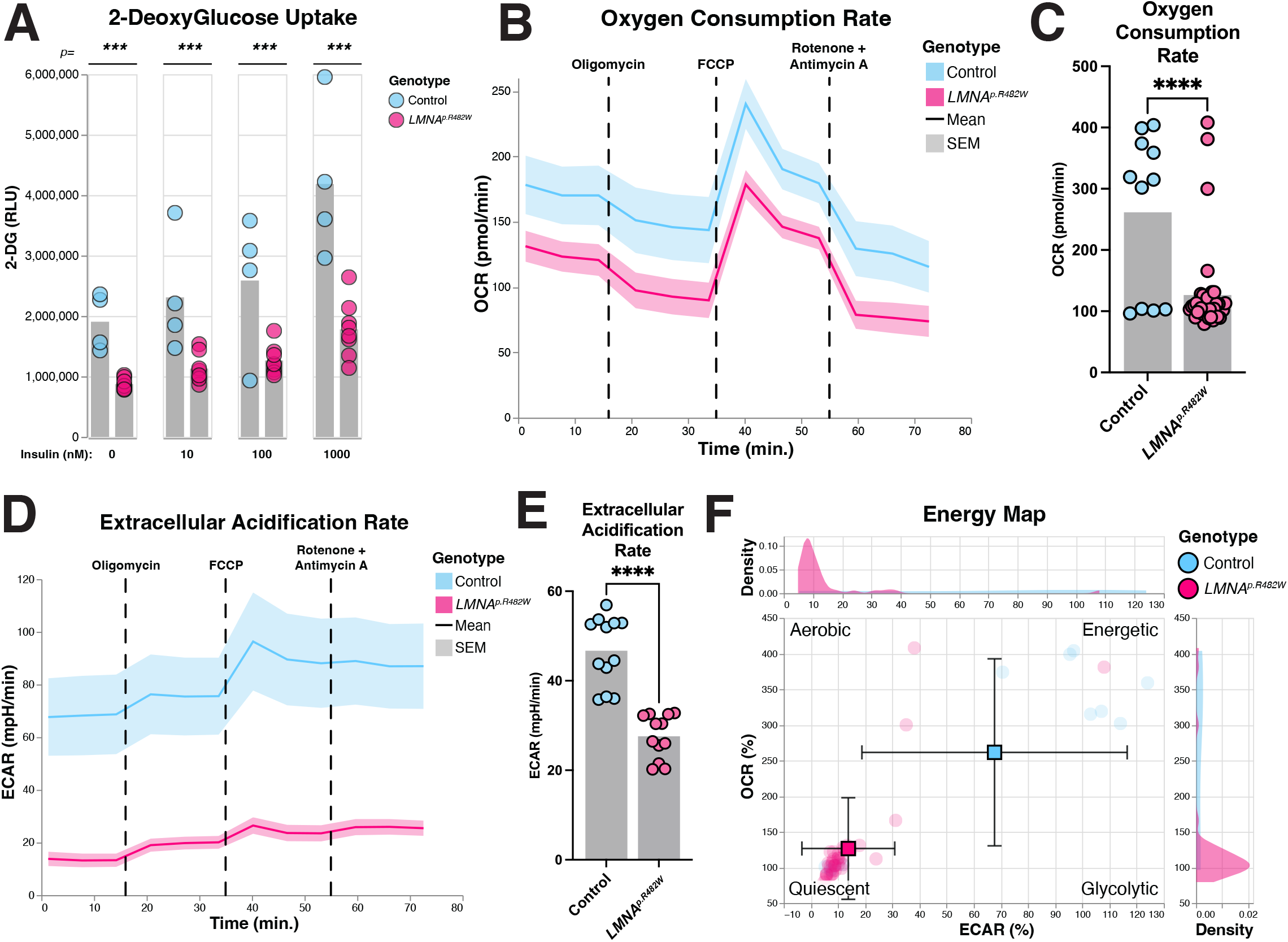
*LMNAp.R482W* iPSC-derived adipocytes exhibit insulin resistance and metabolic quiescence. **(A)** 2-Deoxyglucose (2-DG) uptake in isogenic control (blue) and *LMNA^p.R482W^* (pink) iPSC-derived adipocytes treated with recombinant human insulin at concentrations of 0, 10, 100, and 1000 nM. Y-axis shows 2-DG uptake in relative light units (RLU) normalized to cell viability. Bars represent mean. P-values shown above comparisons are derived from two-tailed unpaired t-tests. **(B)** Representative Seahorse XFe96 oxygen consumption rate (OCR) trace over time for isogenic control (blue) and *LMNA^p.R482W^* (pink) adipocytes. Dashed vertical lines indicate sequential injection of oligomycin, carbonyl cyanide 4-(trifluoromethoxy)phenylhydrazone (FCCP), and rotenone + antimycin A. Line represents mean; shaded area represents SEM. Values normalized to cell number by Hoechst nuclear staining. **(C)** Summary of basal OCR (pmol/min) for isogenic control and *LMNA^p.R482W^* adipocytes. Bars represent mean; points represent individual measurements. **** *p* < 0.0001 by two-tailed unpaired t-test. **(D)** Representative Seahorse XFe96 extracellular acidification rate (ECAR) trace over time for isogenic control (blue) and *LMNA^p.R482W^* (pink) adipocytes. Injection sequence and representation as in panel B. **(E)** Summary of basal ECAR (mpH/min) for isogenic control and *LMNA^p.R482W^* adipocytes. Bars represent mean; points represent individual measurements. **** *p* < 0.0001 by two-tailed unpaired t-test. **(F)** Energy phenotype map plotting ECAR (%) versus OCR (%) for isogenic control (blue) and *LMNA^p.R482W^* (pink) adipocytes. Squares represent group means with SD error bars. Quadrants indicate aerobic, energetic, quiescent, and glycolytic metabolic phenotypes. Marginal density distributions are shown for each axis.

### *LMNA^p.R482W^* adipocytes show reduced oxidative metabolism and metabolic quiescence

To examine the effect of the *LMNA^p.R482W^* variant on mitochondrial function, adipocytes were subjected to treatment with oligomycin, FCCP, rotenone, and antimycin A. Oxygen consumption and extracellular acidification rates were measured using the Seahorse XFe96 Analyzer (Agilent Technologies) and normalized to cell number by Hoechst nuclear staining imaged on a BioTek Cytation 5 at the end of the assay. Overall, the oxygen consumption rate was significantly lower in the *LMNA^p.R482W^* cells compared to isogenic control (WT 261.5 pmol/min vs *LMNA^p.R482W^* 126.5 pmol/min; *p* < 0.0001) (**Fig. 2B-C**; **Fig. S1B-C**). Similarly, the extracellular acidification rate was significantly decreased in the *LMNA^p.R482W^* cells at all time points (WT 46.7 mpH/min vs *LMNA^p.R482W^* 27.6 mpH/min; *p* < 0.0001) (**Fig. 2D-E**; **Fig. S1D-E**). Accordingly, energy mapping of the cell lines using ECAR versus OCR revealed that *LMNA^p.R482W^* cells occupy a markedly quiescent metabolic state compared to isogenic control cells, which demonstrated a more neutral energy balance. *LMNA^p.R482W^* cells exhibited significantly reduced ECAR (13.8 ± 17.1% vs WT 67.6 ± 48.8%; *p* = 3.4 × 10^−7^) and OCR (126.5 ± 71.2% vs WT 261.5 ± 131.2%; *p* = 3.4 × 10^−5^), consistent with global suppression of both glycolytic and oxidative metabolic activity (**Fig. 2F**; **Fig. S1F**).

### WAT biopsies from participants with FPLD2 have decreased fatty acid metabolism and increased inflammatory gene expression

To further validate the iPSC-derived adipocytes, the RNA-sequencing results were compared to snRNA-seq results collected from subcutaneous white adipose tissue biopsies from patients with FPLD2 (6).These participants were previously recruited as part of the LEAD-ABC study. Biopsies were collected from both the upper neck and abdomen from patients with FPLD2 and unaffected control individuals (**Fig. 3A**). For this subset analysis, 3 individuals with the specific *LMNA^p.R482W^* genotype and 5 unaffected controls were studied. Of the 3 individuals with FPLD2, two had developing lipodystrophy and one had established lipodystrophy at the time of biopsy.

**Figure 3.**
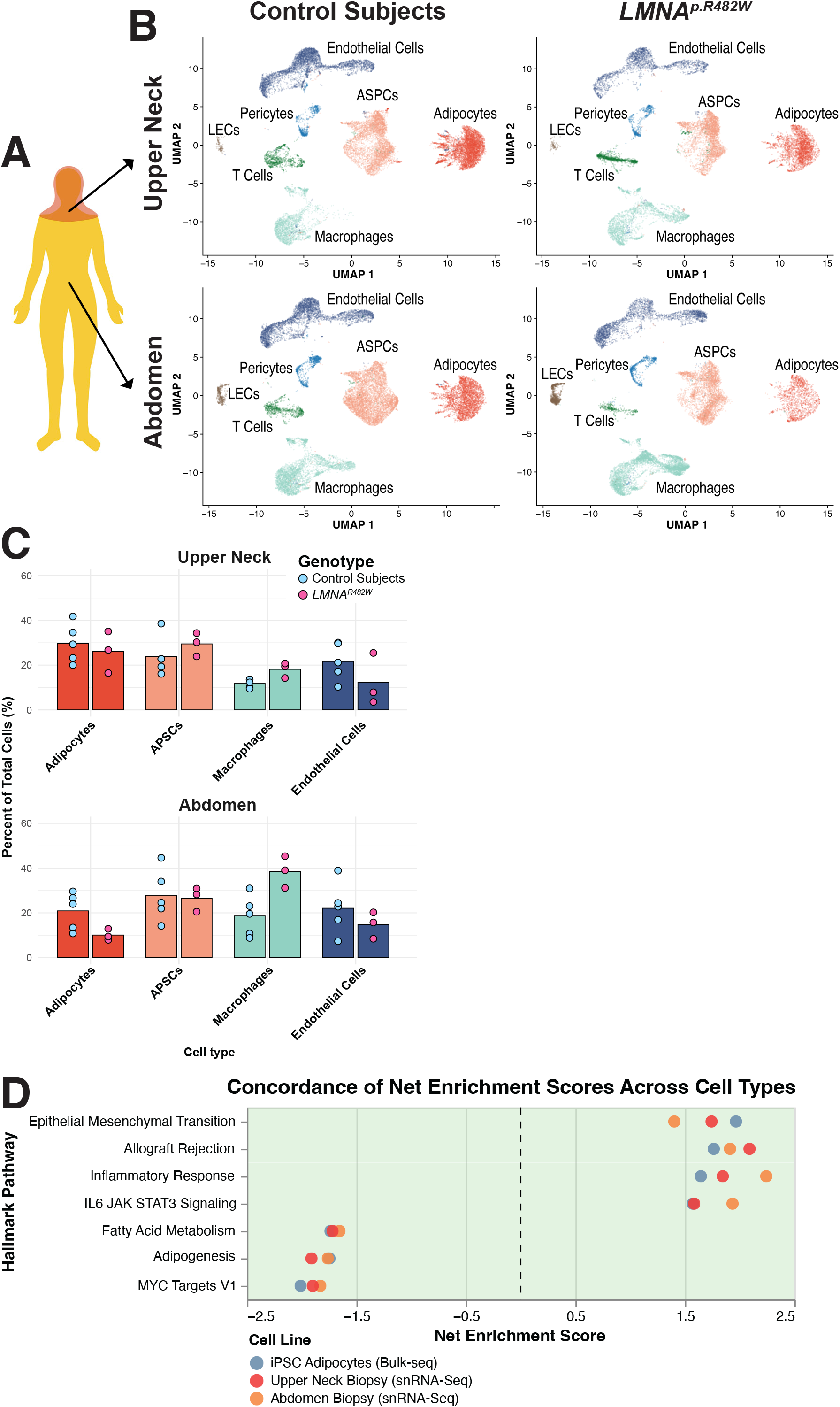
*LMNAp.R482W* iPSC-derived adipocytes recapitulate the transcriptomic signatures of human FPLD2 adipose tissue. **(A)** Schematic of the study design showing anatomical locations of subcutaneous white adipose tissue (WAT) biopsy sites: upper neck (lipohypertrophic depot in FPLD2) and abdomen (lipoatrophic depot in FPLD2). **(B)** Uniform Manifold Approximation and Projection (UMAP) plots of single-nucleus RNA-sequencing (snRNA-seq) data from upper neck (top row) and abdominal (bottom row) WAT biopsies in control individuals (left column) and *LMNA^p.R482W^*individuals (right column). Seven cell populations are labeled: adipocytes, adipose stem and progenitor cells (ASPCs), macrophages, endothelial cells, T cells, pericytes, and lymphatic endothelial cells (LECs). **(C)** Bar charts comparing the percentage of total cells contributed by adipocytes, ASPCs, macrophages, and endothelial cells in upper neck (top) and abdominal (bottom) biopsies between control individuals (open circles) and *LMNA^p.R482W^*individuals (filled circles). Bars represent mean. No differences achieved statistical significance due to the limited sample size. **(D)** Simplified concordance plot of normalized enrichment scores (NES) for seven key Hallmark pathways across iPSC-derived adipocytes (bulk RNA-seq, blue), upper neck biopsy adipocytes (snRNA-seq, red), and abdominal biopsy adipocytes (snRNA-seq, orange). Positive NES values indicate enrichment in isogenic control or control; negative NES values indicate enrichment in *LMNA^p.R482W^*. Pathways shown represent those most central to FPLD2 pathophysiology and were concordantly regulated across all three experimental models.

Seven major cell populations were identified via snRNA-seq of the white adipose tissue biopsies: adipocytes, adipose stem and progenitor cells (ASPCs), macrophages, endothelial cells, T cells, pericytes, and lymphatic endothelial cells (LECs). These cells were independently grouped into clusters, and their identity was assigned via the expression of canonical gene markers (*ADIPOQ*, *PDGFRA*, *MAFB*, *JAM2*, *IL7R*, *STEAP4*, *PROX1*) (**Fig. S4A**) (14).

Uniform Manifold Approximation and Projection (UMAP) was used to plot these 7 clusters across 2-dimensional space, allowing for comparison of the snRNA-seq data between genotype and biopsy location. The most striking results occurred within the abdominal samples, the area of lipoatrophy in individuals with FPLD2. Here, the adipocyte cluster demonstrated a lower density in *LMNA^p.R482W^* individuals compared to controls. Conversely, the macrophage population appeared denser with stronger expression towards UMAP1 in *LMNA^p.R482W^* individuals. Compared to the abdomen, the same UMAP changes for the upper neck samples were present, but less severe (**Fig. 3B**). Cell counts were generated to compare the percentage of total cells that each cluster represented. Due to the small sample size, none of the differences in cell populations achieved statistical significance (**Fig. 3C**).

To explore the broad transcriptomic effects of the pathogenic *LMNA^p.R482W^*variant, GSEA was performed comparing adipocytes from patients with FPLD2 and control individuals. For the upper neck biopsies, the top enriched pathways in control individuals included allograft rejection, inflammatory response, epithelial mesenchymal transition, mitotic spindle, interferon gamma response, apical junction, and IL6/JAK-STAT3 signaling (**Fig. S4B**). There was strong overlap with the abdominal samples, which shared 4 of the 7 most enriched pathways (**Fig. S4C**). The most depleted pathways in *LMNA^p.R482W^* upper neck samples were oxidative phosphorylation, adipogenesis, Myc targets V1, MTORC1 signaling, fatty acid metabolism, protein secretion, and androgen response (**Fig. S4B**). Again, there was strong overlap with the abdominal samples, which shared 6 of the 7 most depleted pathways (**Fig. S4C**).

The pathways most central to FPLD2 pathophysiology, including adipogenesis, fatty acid metabolism, oxidative phosphorylation, and inflammatory response, were concordantly enriched or depleted across iPSC-derived adipocytes, upper neck biopsies, and abdominal biopsies (**Fig. 3D**). To quantify this agreement more broadly, we compared net enrichment scores across all mutually significant Hallmark pathways between the iPSC model and each biopsy site. Against upper neck biopsies, 25 of 37 pathways were correlated in the same direction (Pearson ρ = 0.401; Spearman ρ = 0.532) (**Fig. S5A**), and against abdominal biopsies, 26 of 37 pathways were correlated in the same direction (Pearson ρ = 0.455; Spearman ρ = 0.535) (**Fig. S5B**). In the three-way comparison across all experimental models, 22 of 37 pathways (59.5%) were fully concordant, 7 (18.9%) were partially concordant, and 8 (21.6%) were discordant (Pearson ρ = 0.449; Spearman ρ = 0.529) (**Fig. S6**). These findings strongly support that the isogenic *LMNA^p.R482W^* iPSC system as a faithful *in vitro* model with which to investigate the fundamental principles by which pathogenic lamin A variants alter adipogenic gene expression and disrupt metabolic homeostasis.

## Discussion

In this study, we generated an isogenic iPSC model of familial partial lipodystrophy type 2 (FPLD2) by introducing the pathogenic *LMNA* c.1444C>T (p.R482W) variant into an isogenic control iPSC line using CRISPR/Cas9 genome editing and differentiating these cells to white adipocytes for functional and transcriptomic characterization. *LMNA^p.R482W^* adipocytes demonstrated decreased expression of key adipogenic genes, severe insulin resistance, and global metabolic quiescence. Critically, these *in vitro* phenotypes were validated against single-nuclei RNA-sequencing (snRNA-seq) data from subcutaneous white adipose tissue biopsies of patients with FPLD2, revealing strong concordance of disease-relevant transcriptomic pathways. Taken together, these findings support this iPSC system as a faithful cellular model of FPLD2 adipocyte dysfunction and provide a renewable platform for mechanistic and therapeutic investigation.

The generation of human iPSC-derived adipocytes has emerged as a powerful approach for studying adipose biology and metabolic disease. Foundational work by Ahfeldt and colleagues established that human pluripotent stem cells can be efficiently programmed into white and brown adipocytes via mesenchymal progenitor intermediates, yielding cells with transcriptomic profiles closely resembling primary adipose tissue (7). Building on this foundation, Warren and colleagues demonstrated that iPSC-derived adipocytes from a large population cohort can recapitulate genotype-dependent metabolic variation, supporting their broader utility for functional genomics (8). Su and colleagues further refined the differentiation methodology by generating beige adipocytes from *FOXF1+* mesoderm-derived mural mesenchymal stem cells using commercially defined serum-free conditions, demonstrating that lineage-faithful differentiation without exogenous transgene delivery yields functionally mature adipocytes with robust metabolic activity (16). The present study leverages a similar mesenchymal progenitor-based approach, using MesenCult-ACF adipogenic medium to generate white adipocytes without *PPARG2* overexpression. This design preserves the endogenous transcriptional response to the pathogenic variant in *LMNA* and is an important consideration when contextualizing the findings of the studies described below.

iPSC-based investigation of the pathogenic p.R482W variant in *LMNA* was previously reported by Friesen and Cowan, who reprogrammed fibroblasts from 2 patients with FPLD2 and differentiated them to white adipocytes using lentiviral *PPARG2* overexpression per the protocol described by Ahfeldt and colleagues (9). Their findings of reduced expression of adipogenic markers (*ADIPOQ*, *FABP4*, *HLS*), impaired adiponectin secretion, and attenuated insulin signaling are broadly concordant with the transcriptomic and functional phenotypes described in this study. There are, however, several important methodological distinctions between that study and the present one. Friesen and Cowan compared patient-derived iPSCs to unrelated healthy controls, meaning observed phenotypic differences reflect the combined effect of the *LMNA* variant and inter-individual genetic background variation. In contrast, this study used CRISPR/Cas9 to knock in the p.R482W variant into a single isogenic control background, providing an isogenic comparison that isolates the cell-autonomous effects of the pathogenic *LMNA* variant. Furthermore, lentiviral *PPARG2* overexpression drives adipogenic commitment independently of the endogenous transcriptional program, which may modify or obscure variant-specific effects on adipogenesis. Most strikingly, Friesen and Cowan reported increased maximal oxygen consumption in FPLD2 adipocytes and proposed that elevated lipolysis, which was also increased in their model, liberates free fatty acids as substrate for mitochondrial oxidation (9). In contrast, we observed globally decreased oxygen consumption rate and extracellular acidification rate, with energy mapping placing *LMNA^p.R482W^* cells in a metabolically quiescent state. We propose that this discrepancy reflects the methodological differences described above, in which *PPARG2*-overexpressing adipocytes may have an inherently elevated lipolytic and oxidative baseline that obscures the suppressive effect of the *LMNA* variant on intrinsic metabolic activity. The isogenic design used here, in the absence of forced *PPARG2* expression, may more faithfully unmask the underlying metabolic quiescence of *LMNA^p.R482W^*adipocytes.

The transcriptomic comparison between *LMNA^p.R482W^* and isogenic control adipocytes revealed significant downregulation of multiple genes central to adipocyte identity and function, including *ADIPOQ*, *LIPE*, *ITGA7*, *DHCR7*, *ITIH5*, and *OMD* (13). These genes collectively govern lipid storage and mobilization, cholesterol biosynthesis, integrin-mediated matrix interactions, and extracellular matrix remodeling, processes that are fundamentally disrupted in lipodystrophic adipose tissue. Notably, mesenchymal progenitor cells from the two genotypes showed largely equivalent transcriptomes, with only *ITIH5* significantly reduced in *LMNA^p.R482W^* cells. This stage-specificity suggests that the molecular consequences of the *LMNA* variant emerge primarily during or after adipogenic commitment, consistent with the clinical observation that fat loss in FPLD2 typically manifests at or after puberty rather than being present from birth (2).

The functional demonstration of insulin resistance in *LMNA^p.R482W^*adipocytes using 2-deoxyglucose uptake extends prior work in this area. While Friesen and Cowan inferred insulin resistance from reduced phosphorylation of AKT and ERK1/2 in response to insulin stimulation (9), direct measurement of glucose uptake provides a more physiologically relevant readout of insulin action at the cellular level. The significant reduction in 2-DG uptake observed under both basal and insulin-stimulated conditions across all tested concentrations demonstrates that *LMNA^p.R482W^* adipocytes are intrinsically insulin resistant, independent of systemic factors such as ectopic lipid accumulation or circulating cytokines. This cell-autonomous insulin resistance likely reflects impaired GLUT4 trafficking and downstream insulin receptor signaling (17, 18), and may contribute to the severe metabolic syndrome, including diabetes mellitus and hypertriglyceridemia, observed in patients with FPLD2 (3).

The global metabolic quiescence of *LMNA^p.R482W^* adipocytes, evidenced by concurrent reductions in both oxygen consumption rate and extracellular acidification rate, reflects a broad suppression of both oxidative and glycolytic metabolism. This finding is consistent with the GSEA depletion of oxidative phosphorylation, fatty acid metabolism, and MTORC1 signaling pathways observed in both the iPSC bulk RNA-sequencing data and the human biopsy snRNA-seq data. Suppression of MTORC1 signaling is particularly noteworthy, as this pathway is a key integrator of nutrient sensing, lipid synthesis, and adipogenic differentiation (19). The convergent suppression of multiple energy-generating pathways may reflect a fundamental impairment in the capacity of *LMNA^p.R482W^* adipocytes to establish and maintain normal energy homeostasis, which could contribute to ectopic lipid redistribution and systemic metabolic dysfunction characteristic of FPLD2. An intriguing possibility raised by these findings is that disruption of nuclear lamina integrity in LMNA p.R482W adipocytes impairs the establishment or maintenance of a fully mature metabolic state. Given the established role of lamin A/C in chromatin organization and transcriptional regulation (20), altered nuclear architecture may lead to incomplete activation of adipocyte-specific metabolic programs, resulting in the metabolically quiescent phenotype observed here.

A central strength of this study is the cross-platform validation of the iPSC model against snRNA-seq data from subcutaneous white adipose tissue biopsies of patients with FPLD2 (7). GSEA of the adipocyte cluster in both abdominal and upper neck biopsies revealed enrichment of inflammatory pathways including allograft rejection, IL6/JAK-STAT3 signaling, interferon gamma response, and TNFA signaling via NFκB. Metabolic pathways, including oxidative phosphorylation, fatty acid metabolism, and adipogenesis, were depleted in *LMNA^p.R482W^* individuals. These patterns were recapitulated in the iPSC-derived adipocytes with 59.5% full concordance across 37 mutually significant pathways, with the pathways most central to FPLD2 pathophysiology showing the strongest and most consistent concordance across both systems (14). The relative attenuation of these changes in upper neck compared to abdominal biopsies aligns with the known body fat distribution in FPLD2, wherein the abdomen is a site of progressive lipoatrophy while the neck and supraclavicular regions are relatively spared. These observations validate the iPSC model as a faithful proxy for human FPLD2 adipose tissue and underscore the value of pairing *in vitro* disease models with patient-derived tissue data to establish biological relevance. While rescue experiments (e.g., pharmacologic activation of adipogenic pathways or correction of the *LMNA* variant) could further strengthen causal inference, the isogenic design used here already isolates the effect of this specific *LMNA* variant. Future studies can leverage this platform to systematically evaluate therapeutic strategies aimed at restoring adipocyte function

The model described here carries important translational implications. The convergent transcriptomic and functional phenotypes across iPSC and human tissue models point to several potentially druggable pathways, including PPARγ agonism to restore adipogenic gene expression, mitochondrial support to reverse metabolic quiescence, and anti-inflammatory strategies to attenuate the immune-metabolic crosstalk that characterizes lipodystrophic fat depots. However, human studies using some of these agents in patients with FPLD2 have only been partially effective (21). Beyond therapeutic investigation, the isogenic CRISPR framework provides a scalable platform for the functional characterization of *LMNA* variants of uncertain significance (VUS) in patients with atypical presentations of partial lipodystrophy. Atypical lipodystrophy is increasingly recognized as a clinical entity in which patients present with metabolic features including dyslipidemia, insulin resistance, and abnormal fat distribution, but harbor rare or private variants that fall outside established pathogenic classifications, making diagnosis and clinical management challenging (2). Introducing candidate variants into the same isogenic control AN1.1 background using the approach described here would allow direct side-by-side functional comparisons of adipogenic capacity, insulin sensitivity, and metabolic activity. Such functional evidence could support or refute pathogenicity, potentially improving diagnostic precision and accelerating access to appropriate treatment for patients with lipodystrophy.

Several limitations of the present study warrant acknowledgment. The iPSC model is based on a single isogenic control donor line (AN1.1). While the isogenic design substantially mitigates donor-to-donor genetic confounding, validation across additional genetic backgrounds would strengthen the generalizability of the findings. It must also be noted that, while differentiation efficiency appeared comparable between transgenic and control adipocytes, based on morphology and lipid accumulation, subtle differences cannot be excluded and may contribute to downstream phenotypes. The human biopsy cohort comprised only three individuals with the *LMNA* p.R482W genotype and five unaffected controls, limiting statistical power for cell population-level comparisons. The pathway-level GSEA approach is more robust to small sample sizes and was therefore prioritized for cross-platform comparison. There is an inherent methodological non-comparability between bulk RNA-sequencing of iPSC-derived adipocytes and single-nucleus RNA-sequencing of human biopsies. Accordingly, the use of normalized enrichment scores for pathway-level analysis rather than direct gene-level comparison was intended to bridge this gap. Finally, iPSC-derived adipocytes in monolayer culture lack the paracrine, hormonal, and biomechanical environment of an intact adipose depot, which may limit the recapitulation of phenotypes that depend on cell-cell or cell-matrix interactions *in vivo*. As with all iPSC-derived systems, these adipocytes may not fully recapitulate the maturity of *in vivo* adipocytes, particularly with respect to long-term metabolic adaptation and tissue-level interactions.

In summary, we have generated and characterized an isogenic iPSC-derived adipocyte model of FPLD2 that demonstrates cell-autonomous impairment of adipogenic gene expression, glucose uptake, and mitochondrial metabolism, and that faithfully mirrors the transcriptomic landscape of human FPLD2 adipose tissue. Together, these findings support a model in which LMNA p.R482W drives a state of adipocyte metabolic failure characterized by preserved differentiation but remain functionally immature. This model resolves methodological ambiguities from prior iPSC studies, provides new functional insight into the metabolic quiescence of *LMNA^p.R482W^* adipocytes, and establishes a platform for future mechanistic investigation, therapeutic screening, and the functional classification of *LMNA* variants of uncertain significance in patients with atypical lipodystrophy.

## Author Contributions

SIS, DB, and EAO conceived the studies and planned the experimental design. ABA generated iPSC derived adipocytes. LMA and WB performed imaging studies of iPS derived adipocytes. ABA and MRSG performed functional studies of iPSC adipocytes. MG and DB performed RNA extraction and bulk RNA sequencing of iPSC derived adipocytes. EAO developed the LEAD-ABC study. JNM, AN, and OAM developed the snRNA-seq dataset. WOW and SIS analyzed the bulk RNA-seq and snRNA-seq data. ABA, WOW, and SIS wrote the manuscript, while all other authors edited and approved the final manuscript.

## Conflict of Interest Statement

The authors declare no competing interests.

## Acknowledgement

The authors thank the patients and family members for participating in the LEAD-ABC study whose curiosity of their condition is an everlasting source of purpose.

## Funding and Support

This work is the result of NIH funding, in whole or in part, and is subject to the NIH Public Access Policy. Through acceptance of this federal funding, the NIH has been given a right to make the work publicly available in PubMed Central.

- NIH to SIS (K08 DK124574), OAM and EAO (R01 DK125513), OAM (R01 DK137798, R01 DK121759), JNM (T32 HD007505; F31 DK135181), DB (K08 HG010154).
- Washington University Institute of Clinical and Translational Sciences: Clinical Translational Research Funding Program (CTRFP), ICTS is supported by (UL1TR002345)
- Postbaccalaureate Research Education Program In Diabetes, Endocrinology and Metabolic Diseases (PRIDE; R25DK140763)
- Children’s Discovery Institute - Faculty Recruitment Scholar Award.
- Washington University Division of Physician-Scientists Dean’s Scholar Award.
- Washington University Center for Regenerative Medicine Seed Grant Program.
- Centene/WUSM Precision Medicine Initiative.
- Washington University Genome Engineering Stem Cell Center.
- Washington University Human Cells Tissues and Organoids Center.
- Washington University Diabetes Research Center (P30 DK020579).
- Washington University Nutrition Obesity Research Center (P30 DK056341).
- Washington University Functional Imaging for Variant Elucidation at the McDonnell Genome Institute.
- Michigan Diabetes Research Center (P30 DK020572).
- Michigan Nutrition and Obesity Center (P30 DK089503).
- Caswell Diabetes Institute Atypical Diabetes Program.
- University of Michigan Lipodystrophy Fund (gifted by Sopha, Baker, and Rosenblum families as well as the White Point Foundation of Turkey)

---

**Figure S1.**
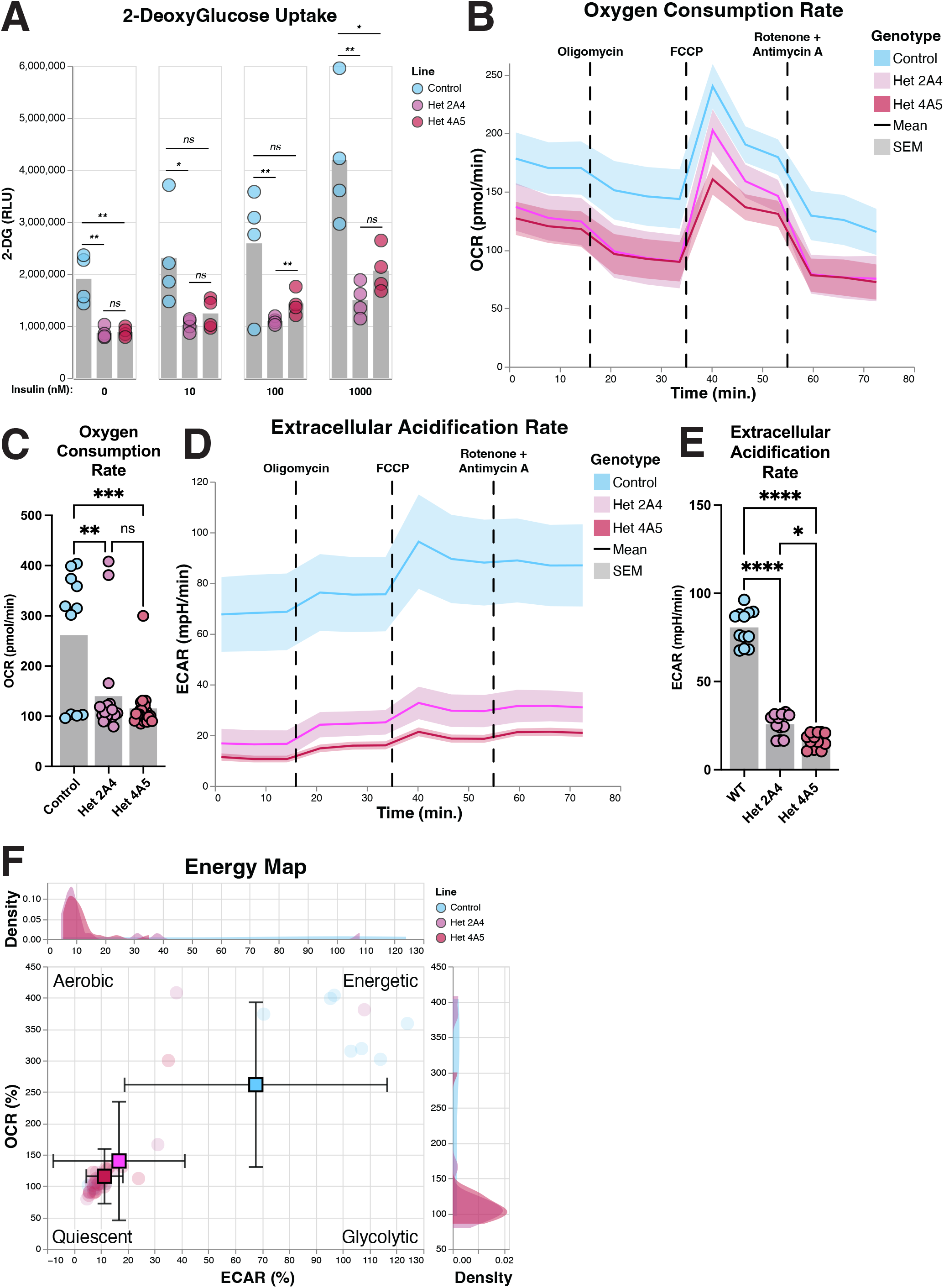
Functional characterization of individual *LMNAp.R482W* iPSC clones. **(A)** 2-Deoxyglucose (2-DG) uptake in isogenic control (blue), Het 2A4 (light pink), and Het 4A5 (dark pink) iPSC-derived adipocytes at insulin concentrations of 0, 10, 100, and 1000 nM. P-values shown above pairwise comparisons. Layout as in Fig. 2A. **(B)** Seahorse XFe96 OCR trace for isogenic control (blue), Het 2A4 (light pink), and Het 4A5 (dark pink) adipocytes over time. Injection sequence and representation as in Fig. 2B. **(C)** Summary of basal OCR (pmol/min) for isogenic control, Het 2A4, and Het 4A5. ** *p* < 0.01; *** *p* < 0.001; ns, not significant, by two-tailed unpaired t-test. **(D)** ECAR trace for isogenic control, Het 2A4, and Het 4A5 adipocytes over time. Layout as in Fig. 2D. **(E)** Summary of basal ECAR (mpH/min) for isogenic control, Het 2A4, and Het 4A5. * *p* < 0.05; **** *p* < 0.0001; ns, not significant, by two-tailed unpaired t-test. **(F)** Energy phenotype map for isogenic control (blue), Het 2A4 (light pink), and Het 4A5 (dark pink) adipocytes. Layout as in Fig. 2F.

**Figure S2.**
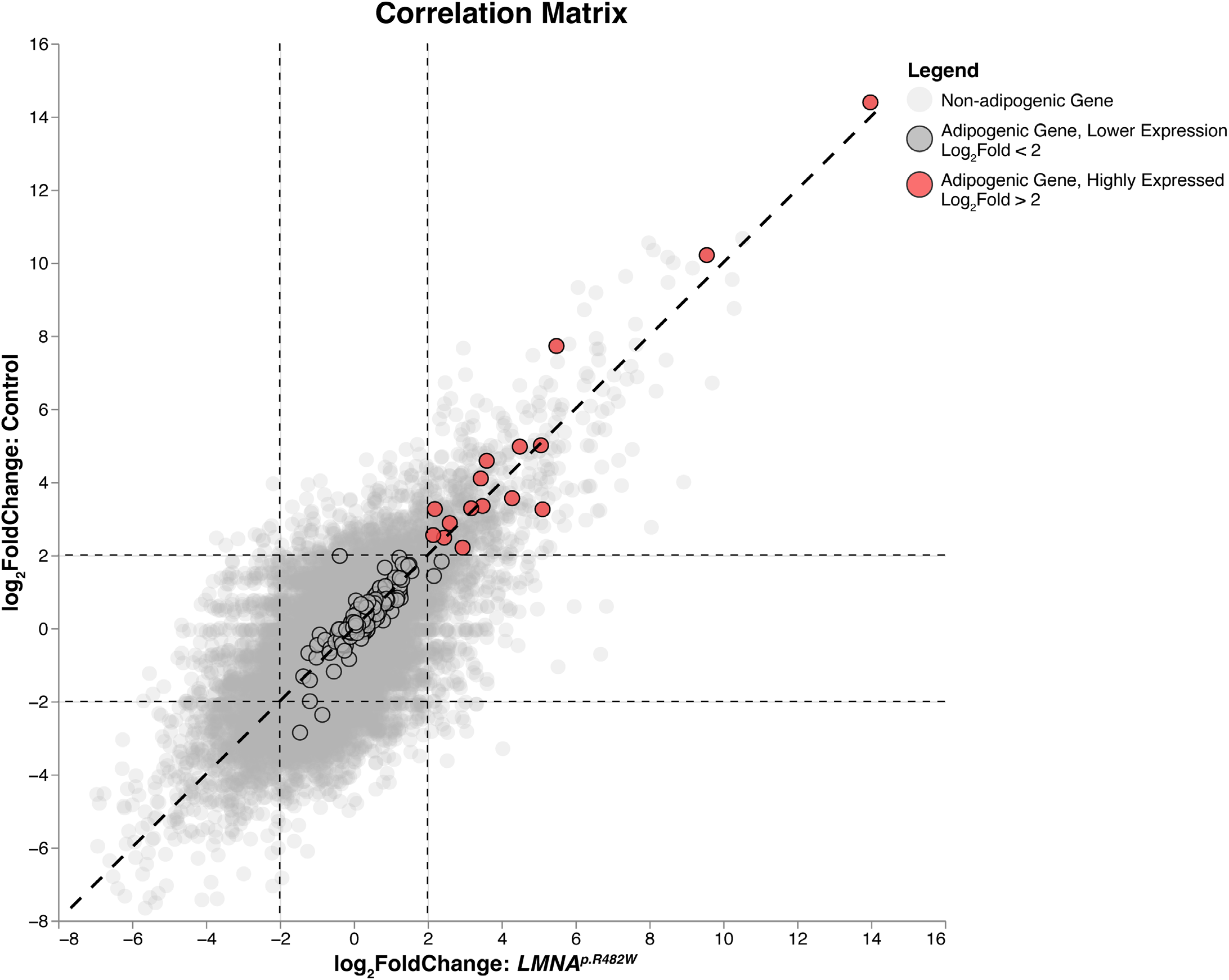
Concordance of adipogenic differentiation gene expression across genotypes. Correlation matrix comparing log_2_ fold changes during adipogenic differentiation (adipocyte vs. MPC) between *LMNA^p.R482W^* (x-axis) and isogenic control (y-axis) cell lines across all expressed genes. Gray points represent non-adipogenic genes; black open circles represent adipogenic genes with log_2_ fold change less than 2; red filled circles represent adipogenic genes with log_2_ fold change greater than 2. Dashed diagonal line indicates perfect concordance. Pearson ρ = 0.700 for all genes; Pearson ρ = 0.968 for adipogenesis gene set.

**Figure S3.**
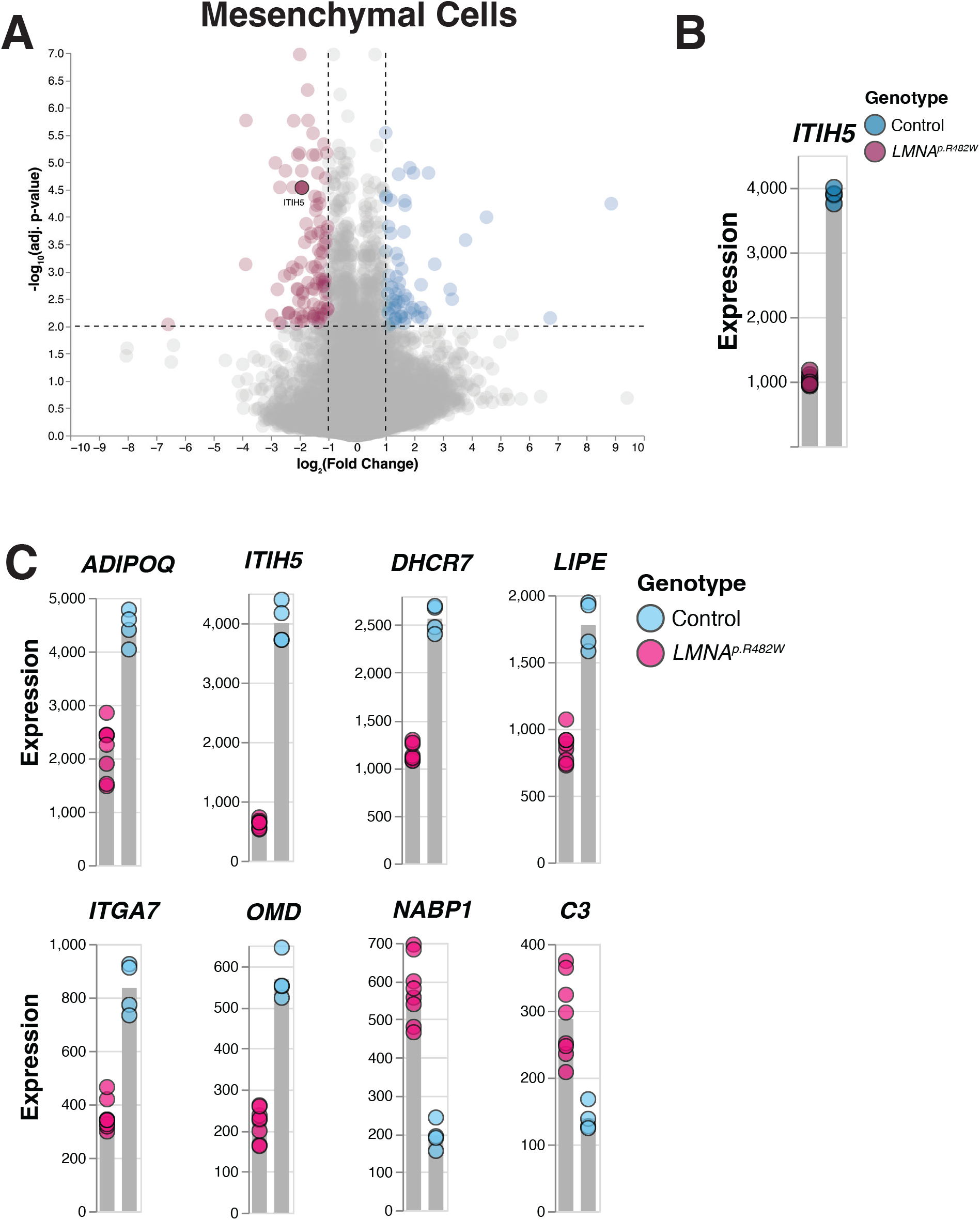
Genotype-specific differences in mesenchymal cells and differentially expressed adipogenesis genes in LMNA^p.R482W^ adipocytes. **(A)** Volcano plot of differential gene expression between *LMNA^p.R482W^* and isogenic control mesenchymal cells. Pink points indicate higher expression in *LMNA^p.R482W^* mesenchymal cells; blue points indicate higher expression in isogenic control mesenchymal cells. *ITIH5* was the only gene achieving statistical significance. Dashed lines indicate log_2_ fold change thresholds of ±1 and adjusted *p*-value threshold of 0.05. **(B)** Expression levels (FPKM) of *ITIH5* in isogenic control (blue) and *LMNA^p.R482W^* (pink) mesenchymal cells. Bars indicate mean. **(C)** Expression levels (FPKM) of the eight significantly differentially expressed Hallmark Adipogenesis genes between *LMNA^p.R482W^* (pink) and isogenic control (blue) adipocytes. *ADIPOQ*, *ITIH5*, *DHCR7*, *LIPE*, *ITGA7*, and *OMD* were significantly lower in *LMNA^p.R482W^* adipocytes; *NABP1* and *C3* were significantly lower in isogenic control adipocytes. Bars indicate mean.

**Figure S4.**
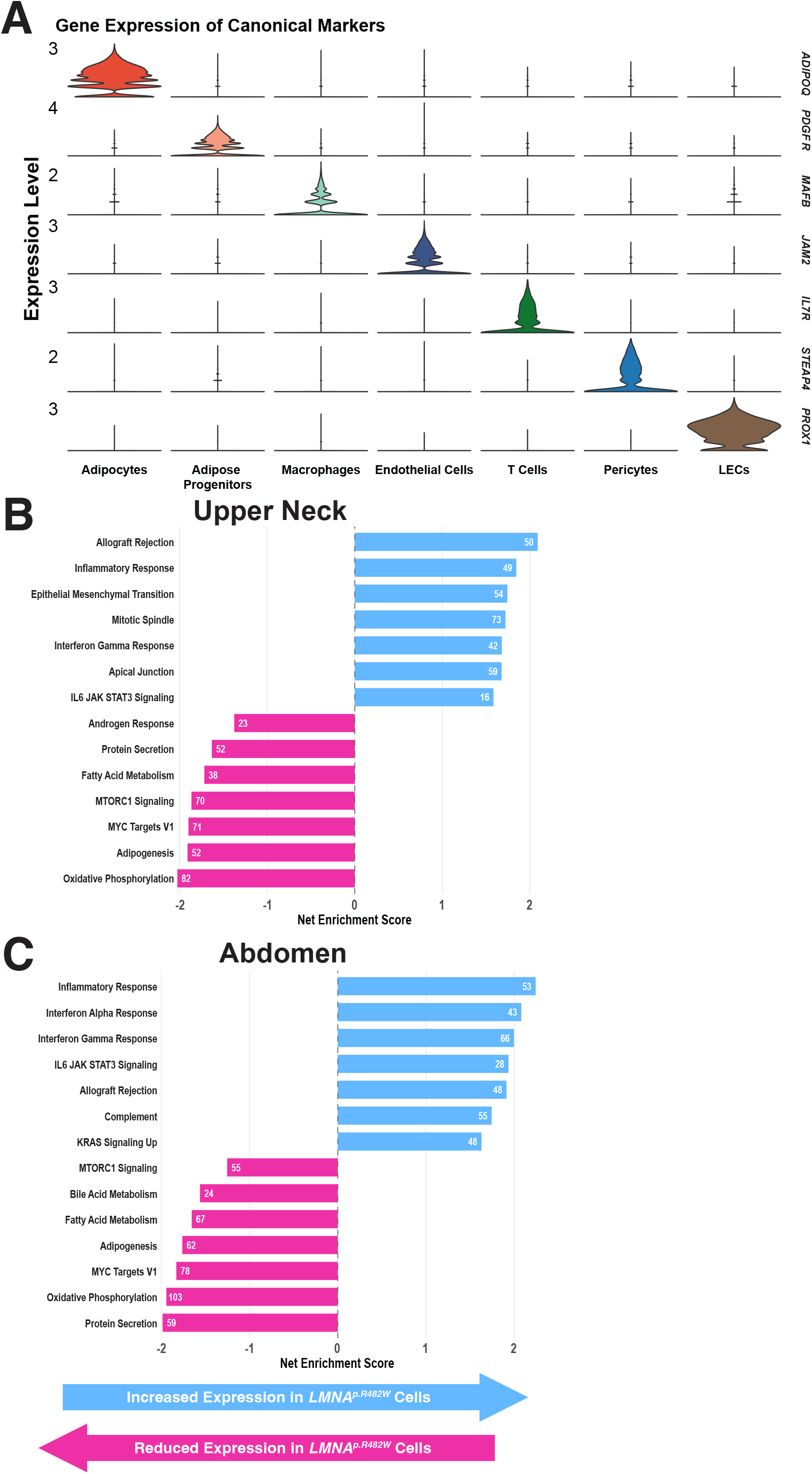
snRNA-seq cell type annotation and GSEA results from human FPLD2 adipose tissue biopsies. **(A)** Violin plots showing normalized expression of canonical cell type marker genes across the seven identified cell populations: *ADIPOQ* (adipocytes), *PDGFR* (adipose stem and progenitor cells), *MAFB* (macrophages), *JAM2* (endothelial cells), *IL7R* (T cells), *STEAP4* (pericytes), and *PROX1* (lymphatic endothelial cells). **(B)** Gene set enrichment analysis (GSEA) bar chart for WAT biopsy-derived upper neck adipocytes comparing *LMNA^p.R482W^*to isogenic control. Blue bars (positive net enrichment score, NES) indicate pathways enriched in *LMNA^p.R482W^* adipocytes; pink bars (negative NES) indicate pathways with reduced expression in *LMNA^p.R482W^*adipocytes. Numbers indicate leading edge gene counts. **(C)** GSEA bar chart for abdominal WAT biopsy-derived adipocytes.

**Figure S5.**
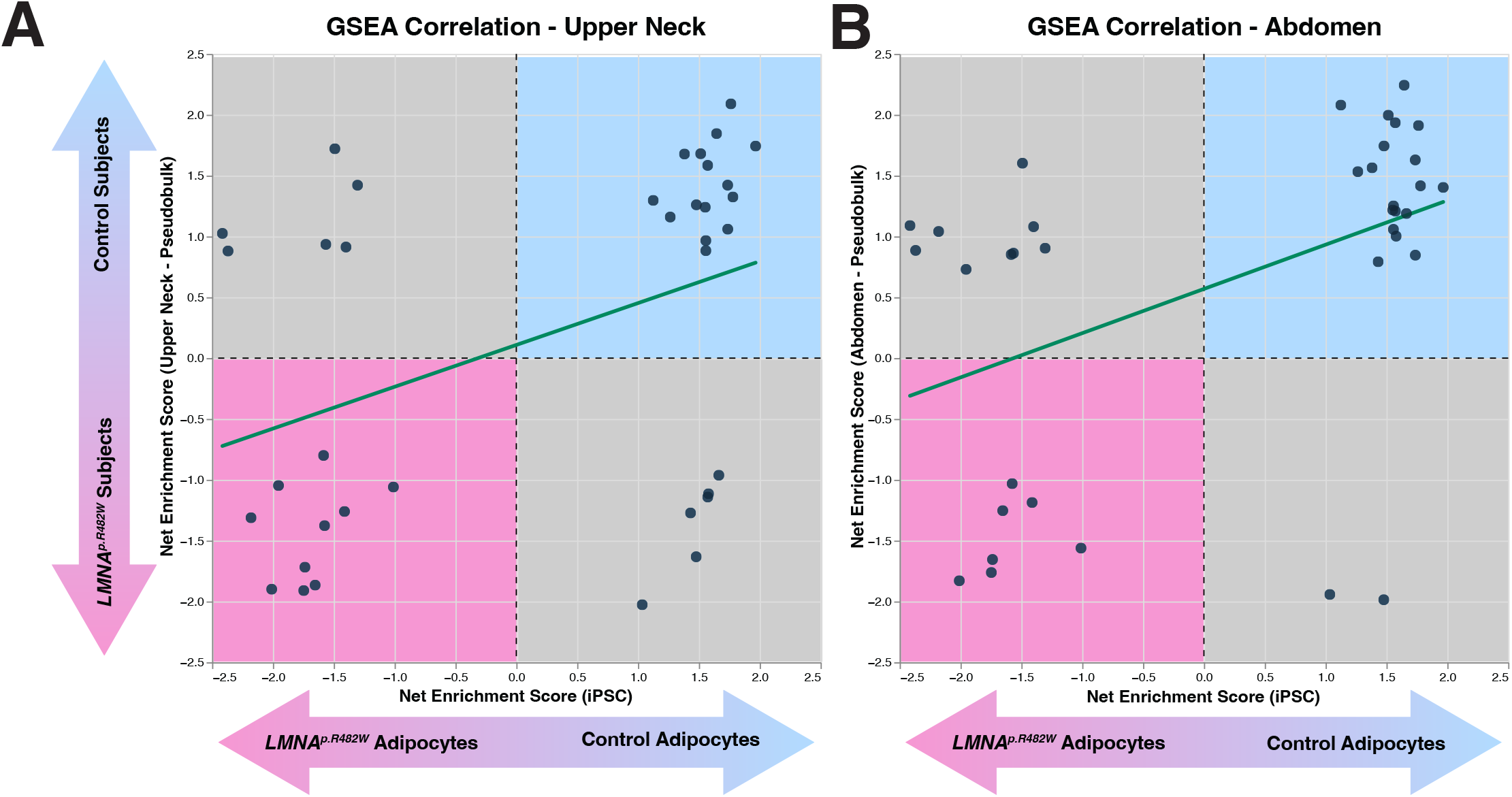
Pairwise correlation of GSEA normalized enrichment scores between iPSC-derived adipocytes and human FPLD2 adipose tissue. **(A)** Scatter plot correlating normalized enrichment scores (NES) from iPSC bulk RNA-seq GSEA (x-axis) and upper neck biopsy snRNA-seq GSEA (y-axis) across mutually significant Hallmark pathways. Positive NES values indicate enrichment in isogenic control or control; negative NES values indicate enrichment in *LMNA^p.R482W^*. Green line indicates linear regression. Pearson ρ = 0.401; Spearman ρ = 0.532; 25 of 37 pathways (67.6%) were correlated in the same direction. **(B)** Scatter plot correlating NES from iPSC bulk RNA-seq GSEA (x-axis) and abdominal biopsy snRNA-seq GSEA (y-axis) across mutually significant Hallmark pathways. Layout as in panel A. Pearson ρ = 0.455; Spearman ρ = 0.535; 26 of 37 pathways (70.3%) were correlated in the same direction.

**Figure S6.**
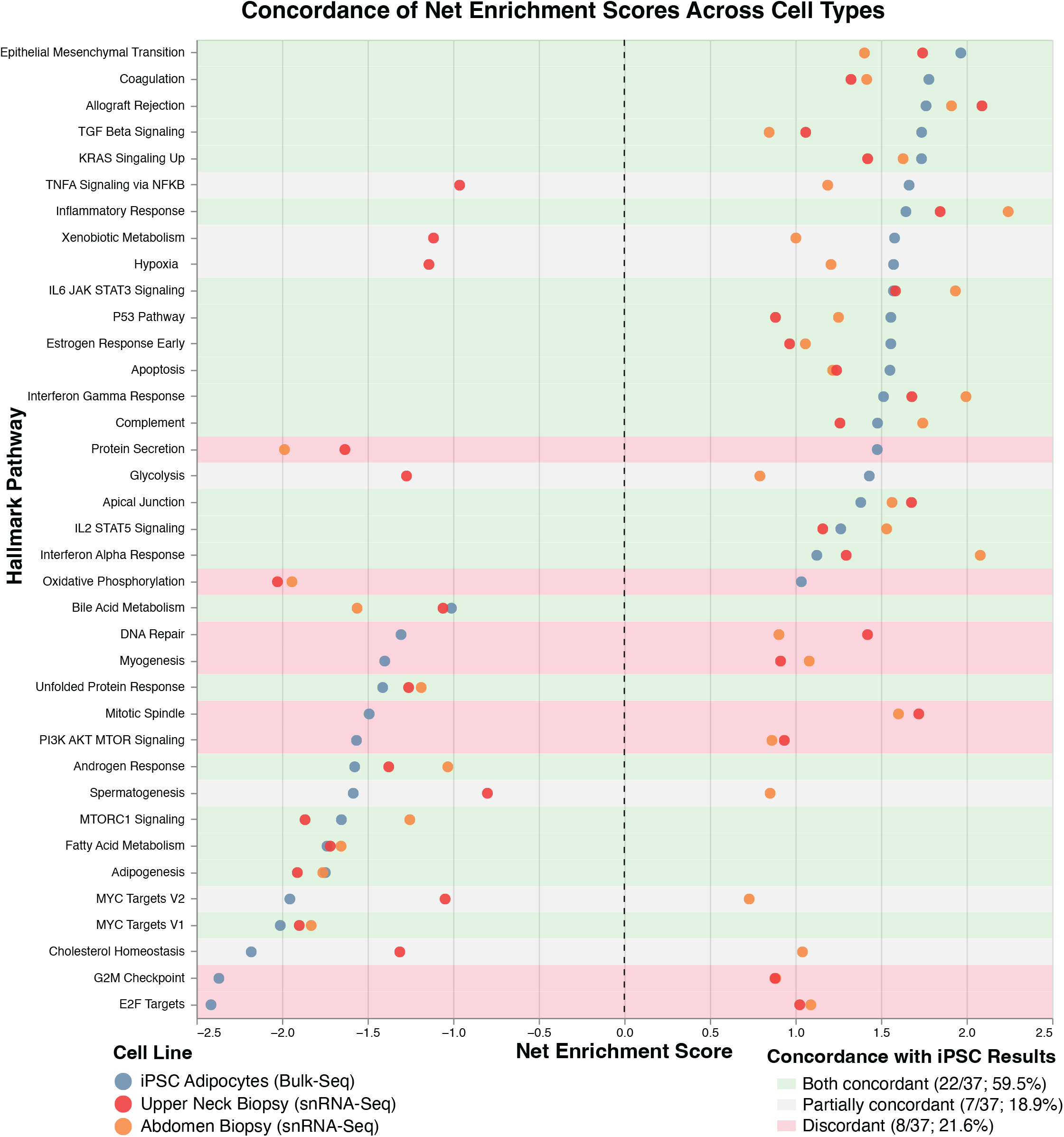
Full three-way concordance of GSEA normalized enrichment scores across iPSC-derived adipocytes and human FPLD2 adipose tissue. Three-way concordance plot of normalized enrichment scores (NES) from GSEA across all 37 mutually significant Hallmark pathways. Each row represents one pathway. Points show NES for iPSC-derived adipocytes (bulk RNA-seq, blue), upper neck biopsy adipocytes (snRNA-seq, red), and abdominal biopsy adipocytes (snRNA-seq, orange). Positive NES indicates enrichment in isogenic control or control; negative NES indicates enrichment in *LMNA^p.R482W^*. Shading indicates concordance classification: green (concordant across all three datasets, 22/37; 59.5%), light green (partially concordant, 7/37; 18.9%), and pink (discordant, 8/37; 21.6%). Leading edge gene counts for control (blue bar, right) and *LMNA^p.R482W^* (pink bar, right) are shown. Pearson ρ = 0.449; Spearman ρ = 0.529.

**Table S1:** Gene Expression of Wild Type Cells.

| Gene Name | Mesenchymal Cells (FPKM) | Adipocytes (FPKM) | *p*- value (adjusted) |
|---|---|---|---|
| *ADIPOQ* | 0.00 | 3983.53 | 1.09E-03 |
| *ANGPT1* | 615.75 | 6252.14 | 3.06E-04 |
| *ANGPTL4* | 340.06 | 3304.06 | 1.76E-03 |
| *APOE* | 468.36 | 11207.66 | 4.10E-04 |
| *CD302* | 22.51 | 131.89 | 1.50E-03 |
| *CD36* | 17.42 | 3705.43 | 2.99E-03 |
| *CIDEA* | 1.08 | 9.78 | 4.76E-03 |
| *FABP4* | 18.85 | 22362.33 | 6.30E-04 |
| *GPX3* | 68.34 | 2188.02 | 3.09E-04 |
| *ITGA7* | 23.82 | 747.26 | 3.02E-03 |
| *LIPE* | 92.89 | 1588.52 | 2.35E-03 |
| *LPL* | 0.00 | 2284.81 | 3.70E-04 |
| *PPARG* | 140.72 | 648.63 | 1.65E-04 |
| *PTGER3* | 3.63 | 33.96 | 1.44E-04 |
| *SORBS1* | 49.12 | 579.98 | 2.36E-03 |
| *STAT5A* | 260.73 | 1917.29 | 6.78E-04 |
| *STOM* | 674.48 | 3752.15 | 3.65E-05 |

**Table S2:** Gene Expression of *LMNA p.R482W* Type Cells.

| **Gene Name** | **Mesenchymal Cells (FPKM)** | **Adipocytes (FPKM)** | ***p*- value (adjusted)** |
|---|---|---|---|
| *ADIPOQ* | 0.13 | 2052.71 | 6.39E-05 |
| *ANGPT1* | 867.22 | 9720.35 | 1.52E-07 |
| *ANGPTL4* | 388.78 | 3520.03 | 2.73E-06 |
| *APOE* | 1086.95 | 13173.24 | 7.73E-04 |
| *CD302* | 32.50 | 144.58 | 1.01E-03 |
| *CD36* | 27.76 | 1246.70 | 1.02E-06 |
| *FABP4* | 19.30 | 14531.15 | 5.10E-06 |
| *FZD4* | 297.09 | 1548.35 | 1.17E-07 |
| *GPX3* | 95.36 | 3195.68 | 4.74E-07 |
| *ITGA7* | 14.95 | 337.30 | 6.65E-06 |
| *LIPE* | 75.01 | 814.48 | 5.17E-06 |
| *PPARG* | 63.62 | 491.46 | 1.50E-06 |
| *PTGER3* | 6.30 | 28.86 | 3.08E-04 |
| *SORBS1* | 24.34 | 475.51 | 9.09E-08 |
| *STAT5A* | 247.18 | 1499.30 | 2.19E-12 |
| *STOM* | 777.28 | 4242.71 | 3.70E-07 |

**Table S3:**
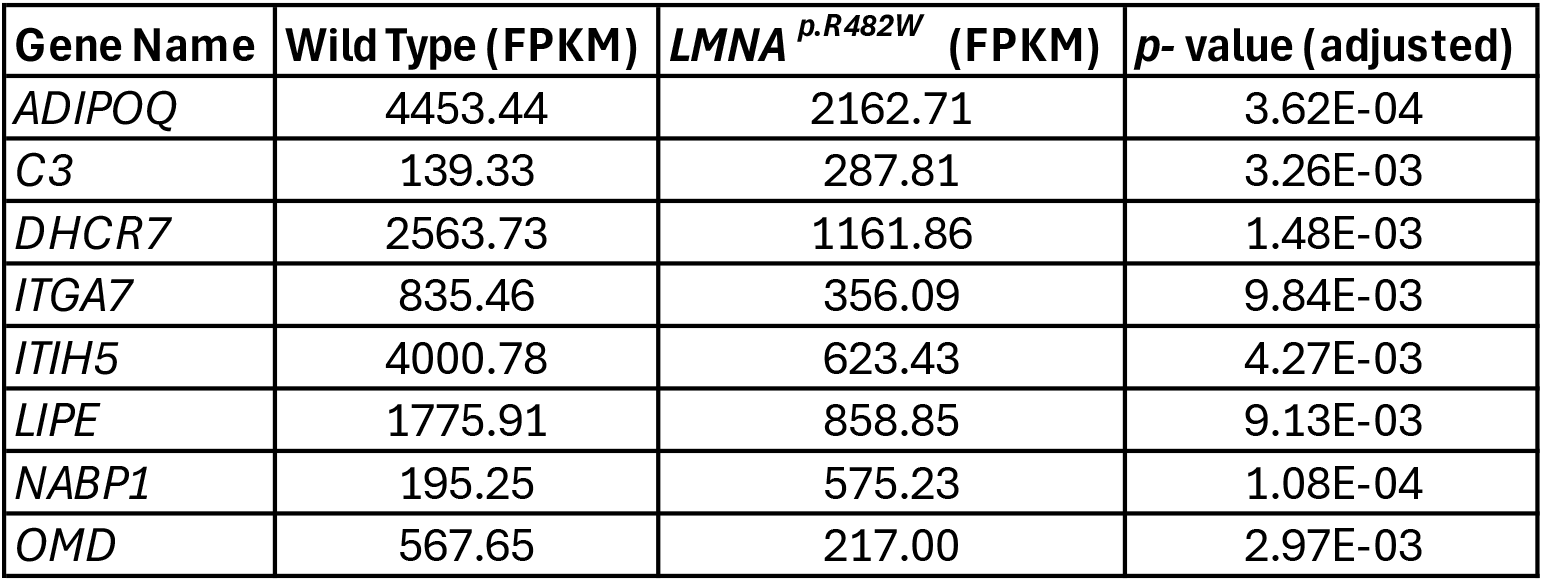
Gene Expression of Adipocytes.

| **Gene Name** | **Wild Type (FPKM)** | **_LMNA_<sup>_p.R482W_</sup> (FPKM)** | **_p_- value (adjusted)** |
|---|---|---|---|
| *ADIPOQ* | 4453.44 | 2162.71 | 3.62E-04 |
| *C3* | 139.33 | 287.81 | 3.26E-03 |
| *DHCR7* | 2563.73 | 1161.86 | 1.48E-03 |
| *ITGA7* | 835.46 | 356.09 | 9.84E-03 |
| *ITIH5* | 4000.78 | 623.43 | 4.27E-03 |
| *LIPE* | 1775.91 | 858.85 | 9.13E-03 |
| *NABP1* | 195.25 | 575.23 | 1.08E-04 |
| *OMD* | 567.65 | 217.00 | 2.97E-03 |

## Notes

### Competing Interest Statement

The authors have declared no competing interest.

## References

1. Lightbourne M, and Brown RJ. Genetics of Lipodystrophy. Endocrinol Metab Clin North Am. 2017;46(2):539–54.

2. Maung JN, Oral EA, and MacDougald OA. Lamin A/C in health, laminopathies, and familial partial lipodystrophy 2. Trends Endocrinol Metab. 2026.

3. Bagias C, Xiarchou A, Bargiota A, and Tigas S. Familial Partial Lipodystrophy (FPLD): Recent Insights. Diabetes Metab Syndr Obes. 2020;13:1531–44.

4. Capanni C, Mattioli E, Columbaro M, Lucarelli E, Parnaik VK, Novelli G, et al. Altered pre-lamin A processing is a common mechanism leading to lipodystrophy. Hum Mol Genet. 2005;14(11):1489–502.

5. Corsa CAS, Walsh CM, Bagchi DP, Foss Freitas MC, Li Z, Hardij J, et al. Adipocyte-Specific Deletion of Lamin A/C Largely Models Human Familial Partial Lipodystrophy Type 2. Diabetes. 2021;70(9):1970–84.

6. Maung JN, Schill RL, Nishii A, Foss de Freitas M, Obua BN, Nygard M, et al. Altered lipid metabolism and inflammatory programs associate with adipocyte loss in familial partial lipodystrophy 2. J Clin Invest. 2026;136(1).

7. Ahfeldt T, Schinzel RT, Lee YK, Hendrickson D, Kaplan A, Lum DH, et al. Programming human pluripotent stem cells into white and brown adipocytes. Nat Cell Biol. 2012;14(2):209–19.

8. Warren CR, O’Sullivan JF, Friesen M, Becker CE, Zhang X, Liu P, et al. Induced Pluripotent Stem Cell Differentiation Enables Functional Validation of GWAS Variants in Metabolic Disease. Cell Stem Cell. 2017;20(4):547–57 e7.

9. Friesen M, and Cowan CA. FPLD2 LMNA mutation R482W dysregulates iPSC-derived adipocyte function and lipid metabolism. Biochem Biophys Res Commun. 2018;495(1):254–60.

10. iPSC line AN1.1 | Washington University Office of Technology Management. https://tech.wustl.edu/tech-summary/ipsc-line-an1-1/.

11. Chen YH, Connelly JP, Florian C, Cui X, and Pruett-Miller SM. Short tandem repeat profiling via next-generation sequencing for cell line authentication. Dis Model Mech. 2023;16(10).

12. Sergushichev AA. An algorithm for fast preranked gene set enrichment analysis using cumulative statistic calculation. bioRxiv. 2016:060012.

13. Liberzon A, Birger C, Thorvaldsdottir H, Ghandi M, Mesirov JP, and Tamayo P. The Molecular Signatures Database (MSigDB) hallmark gene set collection. Cell Syst. 2015;1(6):417–25.

14. Emont MP, Jacobs C, Essene AL, Pant D, Tenen D, Colleluori G, et al. A single-cell atlas of human and mouse white adipose tissue. Nature. 2022;603(7903):926–33.

15. Fernandez-Pombo A, Diaz-Lopez EJ, Castro AI, Sanchez-Iglesias S, Cobelo-Gomez S, Prado-Morana T, et al. Clinical Spectrum of LMNA-Associated Type 2 Familial Partial Lipodystrophy: A Systematic Review. Cells. 2023;12(5).

16. Su S, Guntur AR, Nguyen DC, Fakory SS, Doucette CC, Leech C, et al. A Renewable Source of Human Beige Adipocytes for Development of Therapies to Treat Metabolic Syndrome. Cell Rep. 2018;25(11):3215–28 e9.

17. Bryant NJ, Govers R, and James DE. Regulated transport of the glucose transporter GLUT4. Nat Rev Mol Cell Biol. 2002;3(4):267–77.

18. Govers R. Molecular mechanisms of GLUT4 regulation in adipocytes. Diabetes Metab. 2014;40(6):400–10.

19. Lee PL, Jung SM, and Guertin DA. The Complex Roles of Mechanistic Target of Rapamycin in Adipocytes and Beyond. Trends Endocrinol Metab. 2017;28(5):319–39.

20. Andres V, and Gonzalez JM. Role of A-type lamins in signaling, transcription, and chromatin organization. J Cell Biol. 2009;187(7):945–57.

21. Patni N, Chard C, Araujo-Vilar D, Phillips H, Magee DA, and Akinci B. Diagnosis, treatment and management of lipodystrophy: the physician perspective on the patient journey. Orphanet J Rare Dis. 2024;19(1):263.

